# De novo design of modular protein hydrogels with programmable intra- and extracellular viscoelasticity

**DOI:** 10.1101/2023.06.02.543449

**Authors:** Rubul Mout, Ross C. Bretherton, Justin Decarreau, Sangmin Lee, Natasha I. Edman, Maggie Ahlrichs, Yang Hsia, Danny D. Sahtoe, George Ueda, Nicole Gregorio, Alee Sharma, Rebecca Schulman, Cole A. DeForest, David Baker

## Abstract

Relating the macroscopic properties of protein-based materials to their underlying component microstructure is an outstanding challenge. Here, we exploit computational design to specify the size, flexibility, and valency of *de novo* protein building blocks, as well as the interaction dynamics between them, to investigate how molecular parameters govern the macroscopic viscoelasticity of the resultant protein hydrogels. We construct gel systems from pairs of symmetric protein homo-oligomers, each comprising 2, 5, 24, or 120 individual protein components, that are crosslinked either physically or covalently into idealized step-growth biopolymer networks. Through rheological assessment and molecular dynamics (MD) simulation, we find that the covalent linkage of multifunctional precursors yields hydrogels whose viscoelasticity depends on the crosslink length between the constituent building blocks. In contrast, reversibly crosslinking the homo-oligomeric components with a computationally designed heterodimer results in non-Newtonian biomaterials exhibiting fluid-like properties under rest and low shear, but shear-stiffening solid-like behavior at higher frequencies. Exploiting the unique genetic encodability of these materials, we demonstrate the assembly of protein networks within living mammalian cells and show *via* fluorescence recovery after photobleaching (FRAP) that mechanical properties can be tuned intracellularly, in correlation with matching formulations formed extracellularly. We anticipate that the ability to modularly construct and systematically program the viscoelastic properties of designer protein-based materials could have broad utility in biomedicine, with applications in tissue engineering, therapeutic delivery, and synthetic biology.

**Significance:** Protein-based hydrogels have many applications in cellular engineering and medicine. Most genetically encodable protein hydrogels are made from naturally harvested proteins or protein-polymer hybrid constructs. Here we describe *de novo* protein hydrogels and systematically investigate the impact of microscopic properties of the building blocks (e.g., supramolecular interaction, valencies, geometries, flexibility) on the resultant macroscopic gel mechanics, both intra-and extracellularly. These *de novo* supramolecular protein assemblies, whose properties can be tuned from solid gels to non-Newtonian fluids, provide expanded opportunities for applications in synthetic biology and medicine.

## Introduction

Hydrogels are water-swollen (bio)polymer networks that are widely exploited for bioengineering and medical applications including tissue engineering and drug delivery (1–6). The vast majority of hydrogels explored to date have been created from synthetic polymers [e.g., poly(ethylene glycol), poly(hydroxyethylmethacrylate)] (7–10) or naturally derived proteins (e.g., collagen, fibrin) (11–13). Though such materials have profound utility across many fields, the precursor polydispersity inherent to these systems and a lack of well-defined secondary structure introduces concerns over batch-to-batch variability and imposes challenges in specifying gel attributes *a priori*. Sidestepping many of these limitations, recent efforts have exploited monodisperse recombinant proteins to create hydrogels through covalent and noncovalent interactions (14–17). While these systems have afforded some degree of control over network mechanics (18), prior approaches utilizing native (primarily unstructured) proteins have not permitted systematic investigation of how molecular characteristics of the building blocks (e.g., supramolecular interaction, valencies, geometries, flexibility) influence macroscopic material properties.

*De novo* protein design provides an attractive avenue to computationally specify and vary the molecular characteristics of material building blocks (19). Guided by the physical principles that underlie protein folding, emerging tools such as Rosetta (20) and AlphaFold (21) have made it now possible to design functional proteins with user-specified secondary, tertiary, and quaternary structures. Our group and others have widely applied these techniques to create self-assembling symmetric protein nanomaterials in which the atomic and microscopic properties of the constituents dictate macromolecular structure. These include bounded systems with cyclic (22), dihedral (23), tetrahedral (24), octahedral (25), and icosahedral symmetry consisting of 60 or 120 subunits (26, 27), as well as structurally unbounded 1D fibers (28), 2D arrays (29), and 3D peptide crystals (30), with 3D structures confirmed by x-ray crystallography and cryoelectron microscopy. Though such atomically precise structures have found utility across a wide variety of applications including the rational design of synthetic vaccines (31, 32), modulating cell signaling (33), and molecular motors (34), these methods have not yet been applied towards the creation of bulk materials including hydrogels.

Hypothesizing that *de novo* protein nanostructures could be utilized as branched material crosslinkers with user-specified multivalency, we sought to create homogenous biopolymer gels from these computationally designed protein components and to utilize these perfectly defined materials to isolate and probe the effect of individual microscopic precursor parameters on macroscopic material properties (e.g., viscoelasticity). Exploiting the genetic encodability of this recombinant protein-based approach, we anticipated that such hydrogel precursors could be modularly defined in a plug-and-play manner through conventional cloning alterations of the expression vectors, providing design flexibility for the length, rigidity, density, type, and overall valency of the material crosslinkers. Moreover, since the crosslinking species themselves can be designed with precise structure and symmetry, we anticipated that this strategy would yield more idealized and molecularly homogenous materials than those offered by conventional hydrogel fabrication.

## Results

### Design of *de novo* protein hydrogels with programmable viscoelasticity

We envisioned creating well-defined protein hydrogels through step-growth polymerization methodologies, whereby symmetric homo-oligomers serving as multifunctional network branch points with defined valency would be crosslinked physically or chemically with a reactive protein homodimer (**Figure 1**). Multivalent building blocks were selected from previously *de novo* designed homopentamer (C5, 118.8 kDa as assembled, diameter = 11.3 nm) (23), a 24-chain tetrahedral nanocage (T33, 529 kDa as assembled, diameter = 15.5 nm) (35), and a 120-chain icosahedral nanocage (I53, 1,867.3 kDa as assembled, diameter = 24 nm) (35). To stitch these oligomeric building blocks into bulk gels, we employed a designed homodimer as a divalent material crosslinker with two symmetrical termini pointing in opposite directions (C2, 59 kDa as assembled) (23). C2 and C5 have helical bundles at the core and exterior protruding ‘arms’ that can be readily extended while preserving structural rigidity by the incorporation of additional designed repeat modules (23). Covalent crosslinking of hydrogel precursors (*i.e*., C2 with either C5, T33, or I53) was conducted *via* SpyLigation, a protein “superglue” chemistry, in which an isopeptide bond is irreversibly formed between evolved SpyCatcher-(SC, 88 amino acid residue, 9.5 kDa) and SpyTag-(ST, 13 amino acids) modified species (36). Hydrogel precursors were alternatively crosslinked through the physical association of designed protein heterodimers LHD101-A and LHD101-B (respectively denoted as HA and HB for heterodimers A and B) (37).

**Figure 1:**
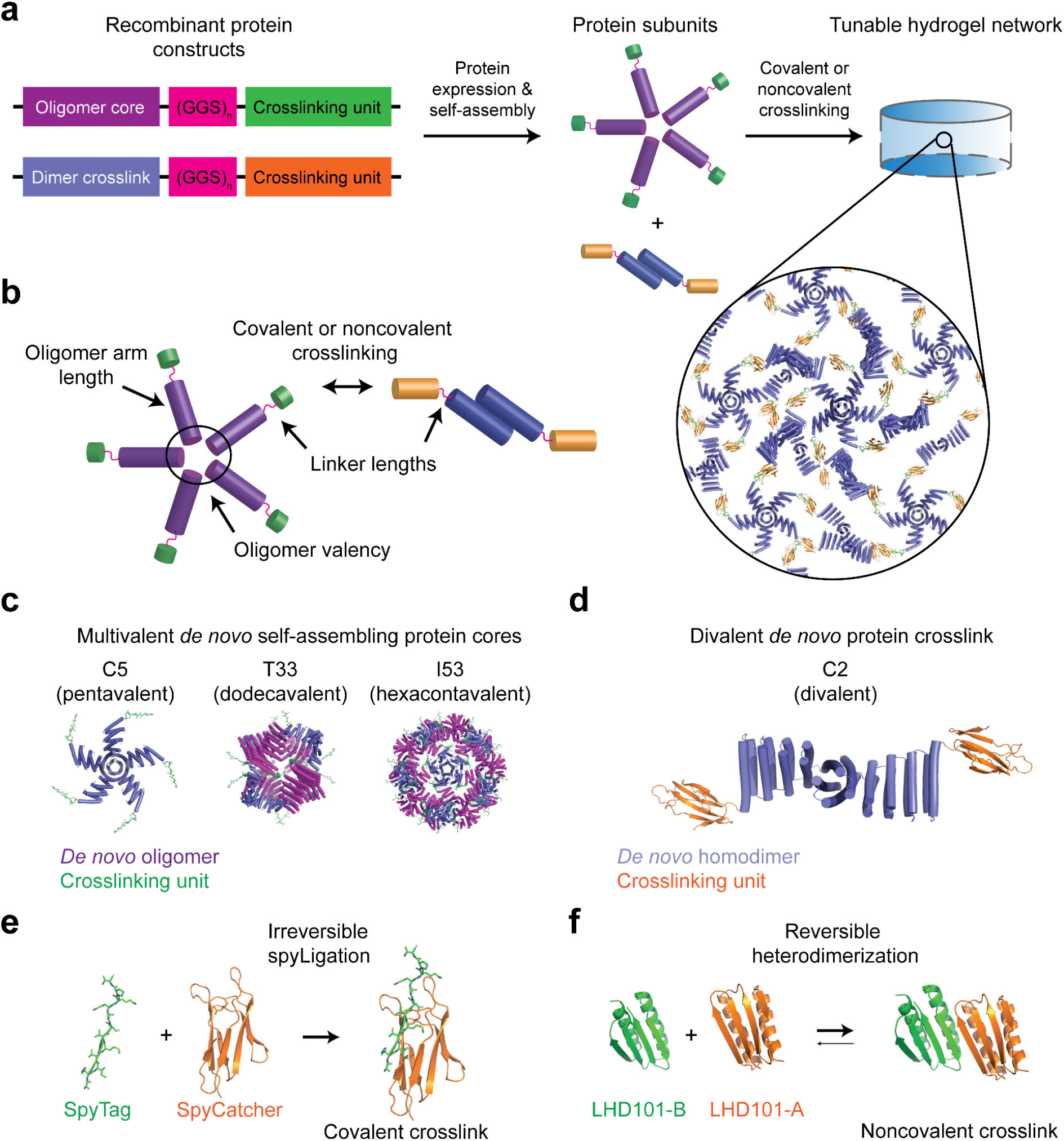
Multi-scale design of *de novo* protein hydrogel networks. a) Protein fusions consisting of self-assembling oligomeric cores fused to crosslinking units are linked by a flexible (GGS)_n_ linker (left) and expressed recombinantly to produce oligomer cores and dimer crosslinks (center), which can then be mixed to form a tunable hydrogel network (right). b) Mechanisms for network tunability explored herein. PDB models depicting three multimeric *de novo* protein cores, showing pendant crosslinking units (here SpyTag) (c); the end-functionalized dimeric protein crosslink (shown here with SpyCatcher) (d); SpyCatcher/SpyTag covalent crosslinking chemistry (e); and the noncovalent crosslinking by reversible heterodimerization of the *de novo* LHD101A/B pair (f).

To create bifunctional *de novo* protein crosslinkers, we first genetically fused each of the self-assembling C2 subunits to SC through a variable-length flexible linker with sequence (Gly-Gly-Ser)_n_ [(GGS)_n_ with n=1 or 5]. C5, T33, and I53 were similarly fused to ST through a (GGS)_n_ linker (n=1, 5, or 10), yielding multivalent self-assembled protein cores that displayed 5, 12, or 60 ST handles, respectively. For the assemblies crosslinked through the physical association of designed protein heterodimers, only the C2-(GGS)_5_-HA: C5-(GGS)_5_-HB pair was chosen. For clarity, we adopted a standard naming convention referring to the 13 resultant protein complexes: protein assemblies are denoted “X-Y-Z”, where “X” is the underlying multivalent protein core, “Y” is the number of GGS repeats in the flexible linker, and “Z” is the crosslinking functional group (e.g., I53-5-ST refers to the I53 core with 5 GGS repeats between it and each of the 60 total ST functional groups). Following plasmid construction through standard cloning techniques, 6xHis-tagged proteins were expressed recombinantly in Lemo *E. coli*, solubly purified by Ni-NTA immobilized-metal affinity chromatography, buffer exchanged into tris-buffered saline (300 mM NaCl, 25 mM Tris, pH 8.0), and concentrated as self-assembled complexes to 100-300 mg/mL (23). Protein purity was confirmed by SDS-PAGE analysis.

Bulk hydrogels were formed by mixing multimeric protein components to a fixed final concentration (10% w/v) with equal stoichiometry between reactive end groups (*i.e.*, SC and ST, HA and HB). Solution viscosity noticeably increased within seconds of component mixing, consistent with rapid SpyLigation kinetics (36) and large heterodimer binding affinities (37). To ensure complete gelation, network formation was allowed to proceed at room temperature (25 ^0^C) for >6 h and analyzed by SDS-PAGE (**Figure S1**) prior to bulk materials characterization. Free-standing macroscopic gels were obtained for all tested combinations.

### Linker flexibility

To assess how the flexibility of the linkers connecting the components affect material viscoelasticity, we measured the rheological properties of gels made from different combinations of C2 × C5 with different length flexible linkers between the rigid homo-oligomers and the covalent SC/ST crosslink (**Figure 2a**). Representative images of hydrogels formed 12 hours after mixing the components are shown in **Figure 2b**. 0.5mm-thick hydrogel droplets with an 8mm diameter were subjected to frequency sweeps between parallel plates ranging from 0.1 to 500 rad/s at a constant strain of 10%, which was empirically determined to lie within the linear viscoelastic regime of these hydrogels. The stiffness of the hydrogel (measured through storage modulus within the linear viscoelastic region) gradually increased as the length of the (GGS)_n_ linker increased (**Figure 2c**).

**Figure 2:**
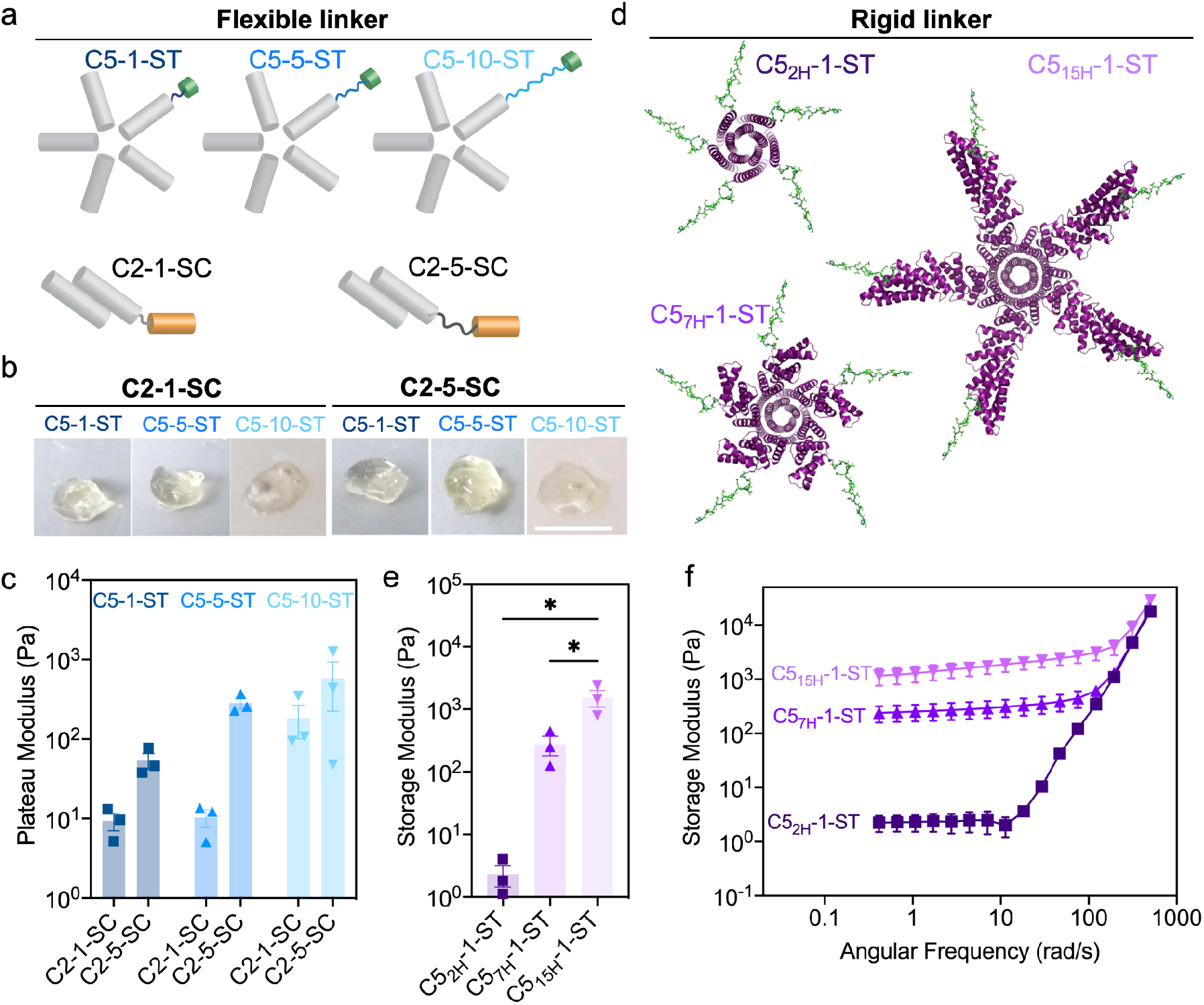
Influence of linker length and rigidity on covalent hydrogel viscoelasticity. The mechanical properties of the hydrogels are dictated by the lengths of the linker connecting the interacting domains. a-c) Flexible linkers. a) Schematics showing C5 and C2 helical bundle oligomers with variable (GGS)_n_ linker. b) Image of hydrogels formed by C2-1-SC and C2-5-SC with ST (SpyTagged) C5 (*i.e.,* C5-1-ST; C5-5-ST; and C5-10-ST]. Scale bar 4 mm. c) Storage modulus of various C2 × C5 combinations. d-f) Rigid connecting elements. d) PDB models of C5 oligomers with 2, 7, and 15 helical lengths. e) Storage modulus of C2-1-SC × C5_nH_-1-ST (rigid arm) combinations derived from angular frequency sweep (f). Dashed lines indicate ± SEM. *p < 0.05 by one-way ANOVA and Holm-Sidak post hoc test.

### Rigid arm length

We next investigated how the size of structured hydrogel components affect the rheological properties of the gels. We took advantage of the ability to computationally vary the length of the rigid helical repeat arms extending from the helical bundle central cores. Each repeat, comprised of two helices (38), is encoded by an identical sequence of 42 amino acids, and the length of the rigid arms can be varied modularly simply by inserting or deleting sequence repeats. Keeping the flexible interaction domain (SC/ST) at a minimum distance of (GGS)_1_, we generated a series of pentameric protein complexes with arm lengths of 2 helices (C5_2H_-1-ST, only the core pentameric helical bundle without extra repeat extension), 7 helices (C5_7H_-1-ST), and 15 helices (C5_15H_-1-ST), respectively (**Figure 2d**), and again measured gel mechanics through frequency sweep rheology. Note that the rigid arm length from the core varies accordingly depending on the repeat extension: ∼1 nm for C5_2H_; ∼3.7 nm for C5_7H_; and ∼7.5 nm for C5_15H_. Similar to the results with varying flexible linker length, we observed a steep increase in the storage modulus as the rigid arm length increased from 1.0 to 7.5 nm (**Figure 2e**). Networks formed with longer rigid arms also exhibited broader viscoelastic regions in the frequency domain: gels with 2H crossed into glassy behavior at an angular frequency around 10 rad/s, while increasing the number of rigid repeats to 15H increased this crossover by nearly a full order of magnitude (**Figure 2f**).

### Valency

We next investigated the effect of building block valency on hydrogel properties. We replaced the C5 SpyCatcher tagged component with one of two two-component protein nanocages – a tetrahedral (T33 symmetry, 24 chains, 12 ST) and an icosahedral (I53 symmetry, with 120 chains and 60 ST). To assemble hydrogels, these nanocage-STs were mixed with C2-SC with different (GGS)_n_ linker lengths. Gel formation was observed for all combinations after mixing, but with starkly different kinetics. The C2 × I53 combinations exhibited rapid gel formation in minutes, while the C2 × T33 combinations formed gels on the hour time scale (**Figure 3a**; 40 microliters of each combination after 12 hours were imaged using a regular camera). Rheological measurement of the elastic moduli revealed that, for both C2 × T33 and C2 × I53 combinations, the storage modulus dropped as the (GGS)_n_ linker length increased (**Figure 3b-c**), in contrast to the results with the smaller C5 core, where stiffness increased as the linker length increased (**Figure 2c**). In networks formed with higher-valency structured components, gel rigidity may no longer be limited by the extent of the SC/ST reaction; instead, increasing linker length likely reduces network stiffness by giving more flexibility to regions between the large oligomeric protein cores.

**Figure 3.**
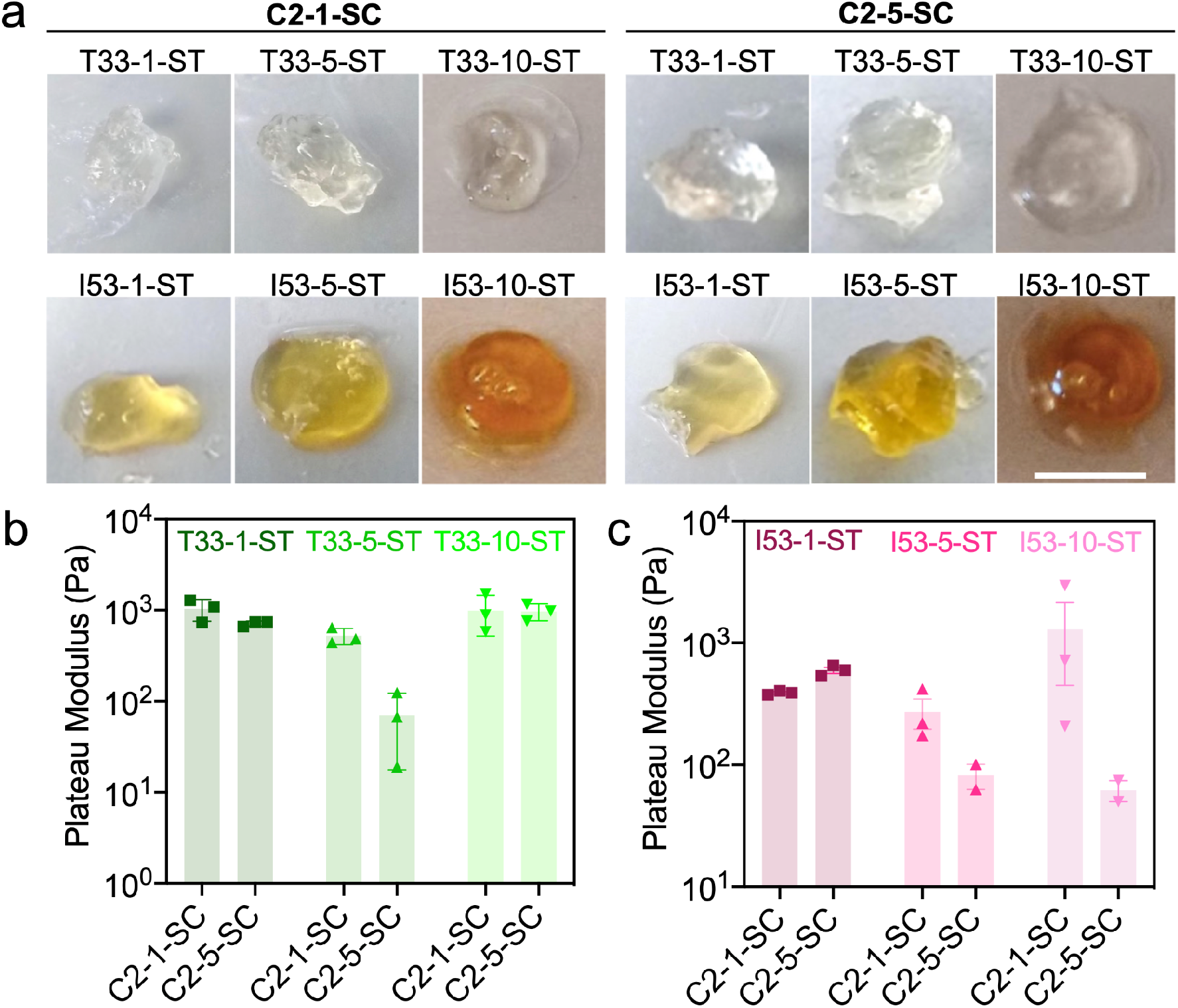
Effect of component valency on hydrogel properties. The elastic modulus of the hydrogels made from higher valency protein assemblies generally decreased as the length of the linker connecting the interacting domains of C2 increased, unlike in the C5 case (Figure 2c). a) Gels formed by C2 × T33 and C2 × I53 combinations. Scale bar 4 mm. b-c) Storage moduli of combinations of C2 × T33 (b) or C2 × I53 (c) with different flexible linker lengths.

To relate the rheological measurements to building block properties at the molecular level, we turned to molecular dynamics (MD) simulations. We developed a simplified protein model for the C2 × C5 systems with varying linker lengths using the HOOMD-Blue MD simulation toolkit (39), inspired by a coarse-grained model of DNA-grafted colloids (40). In this model, C2, C5, and SC are represented by rigid bodies of spheres connected by flexible (GGS)_n_ linkers and ST (**Figure S2**). To evaluate the porosity of the gel as a function of linker length, we calculated the local density fluctuations of the networks on a regular grid (**Figure 4a-b**). We found that the shortest linker gels had the highest porosity, with the fraction of empty grid voxels decreasing as the linker length increased (**Figure 4d**). This linker length-dependent porosity reflects the local clustering of the gels: the intensity of the first peak of the radial distribution function g(r) (**Figure 4c and 4e**) is stronger for the shorter linker gels indicating higher local clustering (41). The longer linker gels have radial distribution functions with less pronounced peaks, indicating more isotropic gel networks with fewer voids (**Figure 4f**). These results are consistent with our experimental observations: shorter linkers lead to more porous networks with lower stiffness.

**Figure 4:**
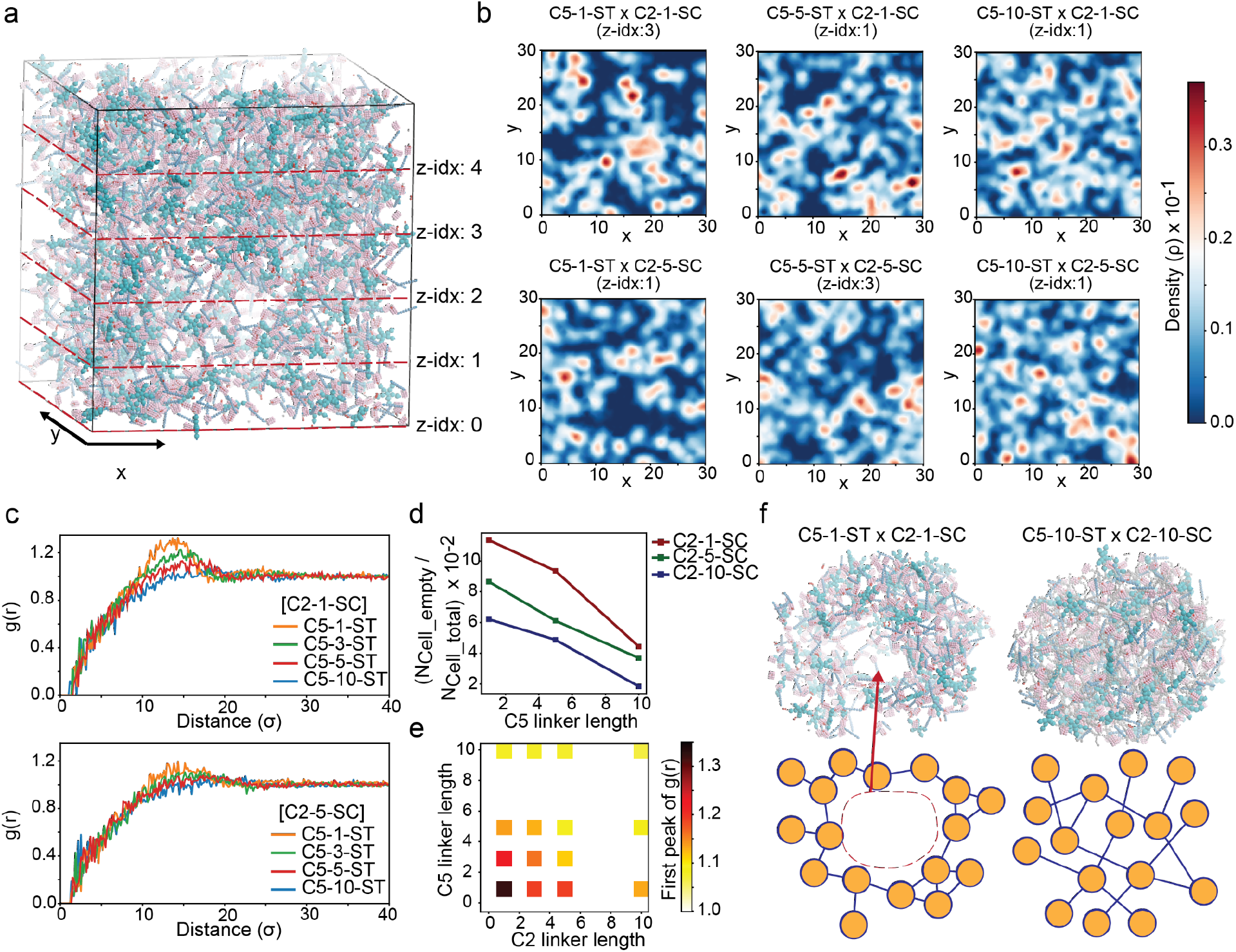
Molecular dynamics (MD) simulation of covalently cross-linked *de novo* protein hydrogels. a) A system of C5-1-ST and C2-1-SC forming a gel network under isochoric conditions after MD equilibration. b) Density distributions of cross-sections of the simulated gel networks from different C2 × C5 combinations. c, e) Radial distribution function plots and the intensity of the first peak of different C2 × C5 combinations. d) Fraction of empty grid voxels for the simulated gels. Voxel dimensions are × × y × z = 3000×3000×5. f) Local structure of (left) C2-1-SC × C5-1-ST and (right) C2-10-SC × C5-10-ST systems.

### Cytocompatability

Having demonstrated that modulate de novo protein-based hydrogels can be formed with tunable viscoelasticity from well-defined macromolecular precursors, we next investigated whether these systems would permit cytocompatible cell encapsulation and sustained 3D cell culture. As an initial test, 10T1/2 fibroblasts were suspended in a phosphate-buffered saline (PBS) solution of C2-5-SC prior to mixing with C5-5-ST. Following polymerization (10 µL droplets) at 37°C, cells were maintained in culture for 24 hours, with viability assessed. High viability was observed in each case (**Figure S3**), indicating the suitability of *de novo* protein-based hydrogels for 3D cell culture and its potential utility in tissue engineering.

## Noncovalent protein networks exhibit non-Newtonian properties both intra- and extracellularly

We next investigated the impact on material viscoelasticity when replacing the covalent interaction in the C2 × C5 hydrogel with a non-covalent, and thus reversible, interaction (**Figure 5a**). We swapped the SC/ST with two chains of the *de novo* designed heterodimer LHD101, henceforth referred to as HA and HB, which are ∼8.6 kDa each and interact with nanomolar binding affinity with on/off rates of ∼2×10^6^ M^-1^ s^-1^ and ∼4×10^-3^ s^-1^ respectively (37). C2-5-HA was mixed with C5-5-HB at equimolar ratios of HA and HB, and the mixture was inspected for viscoelastic material formation. The noncovalent mixture appeared droplet-like (**Figure 5b**, bottom left), in comparison to the analogous system with covalent SpyLigation-based interactions (**Figure 5b**, top left). 10 minutes following the application of a stress, the deformed noncovalent material reverted back to a droplet-like fluid, whereas the covalent materials remained as distorted gels (**Figure 5b**). Further rheometric characterization showed that the noncovalent materials exhibited a lower storage modulus than loss modulus (G’<G”) at resting or low frequency indicating liquid-like properties; the storage moduli overtook the loss moduli (G’>G”) at higher frequencies. In contrast, the covalent gels invariably showed a higher storage modulus than the loss modulus (G’>G”) throughout the frequency sweep, indicating that these materials persisted throughout as solid gels (**Figure 5c**). Given the noncovalent nature of the material crosslinks, we hypothesized that LHD101 gel networks would exhibit self-healing characteristics, which was verified by cyclic strain time sweep rheology (42): the LHD101 gel networks were capable of self-healing over multiple cycles of 500% strain (**Figure S4**).

**Figure 5:**
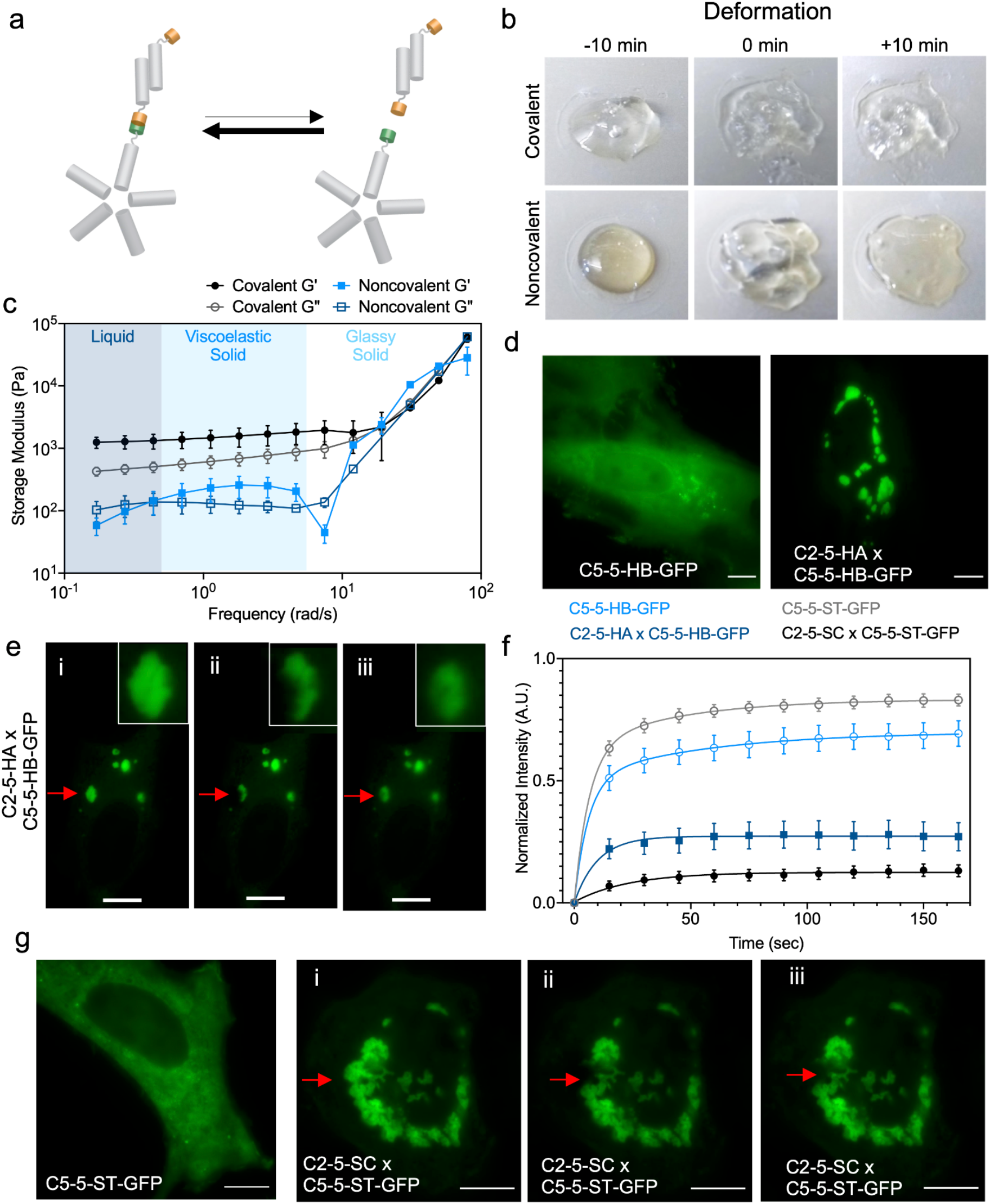
Non-Newtonian fluid properties of the non-covalent assemblies. a) Schematics showing the ON/OFF equilibrium of the LHD101 heterodimer within the C2-5-HA × C5-5-HB protein network. b) Comparison of covalent vs noncovalent hydrogels through deformation studies [Covalent: C2-5-SC × C5-5-ST; Noncovalent: C2-5-HA × C5-5-HB]. Pictures were taken 10 mins before, at 0 mins (at the point of deformation), and 10 mins after deformation. Scale bar 4 mm. c) Rheometric studies (frequency sweep) show that the noncovalent assemblies behave as a viscoelastic liquid at resting or low frequency (G’<G”), but exhibit gel-like (viscoelastic solid) properties as the frequency is increased (G’>G’’). In contrast, covalent gels behave as viscoelastic gels throughout the frequency sweep (G’>G’’). At very high frequency, both covalent and noncovalent assemblies behave as glassy solids. d) Designed non-covalent assemblies form in cells. C5-5-HB-GFP alone (left panel), and coexpression of C2-5-HA and C5-5-HB-GFP (right panel). e) FRAP experiment showing intracellular noncovalent C2-5-HA × C5-5-HB-GFP assemblies as non-Newtonian fluid droplets, similar to the extracellular experiment. (i) before photobleaching, (ii) immediately after photobleaching, and (iii) after complete recovery (180 seconds). f) Time-dependent fluorescence recovery plot of noncovalent assembly (C5-5-HB-GFP, control; and C2-5-HA × C5-5-HB-GFP), and covalent assembly (C5-5-ST-GFP, control; and C2-5-SC × C5-5-ST-GFP). Error bars indicate ± SEM. g) Designed covalent gels formed in cells. Expression of C5-5-ST-GFP in cells alone showed diffuse fluorescence, whereas coexpression of C5-5-ST-GFP and C2-5-SC led to the formation of stable intracellular punctae. FRAP experiments showing (i) before photobleaching, (ii) immediately after photobleaching, and (iii) after 180 seconds. Scale bar 10 µm.

This soft fluid behavior at resting or at low applied frequency but gel-like behavior at a higher frequency is characteristic of non-Newtonian shear thickening (43). Analogous non-Newtonian fluid properties in native protein materials are rare to our knowledge, though certain biopolymers, such as oobleck (cornstarch in water), exhibit similar non-Newtonian behavior (44). We hypothesize that rapid equilibration between LHD101 heterodimers endows the material with fluid-like properties at the resting state, but once the stress is applied at a frequency faster than the on and off rates of the LHD101 A/B interaction, the intermolecular interaction between C2 and C5 (through HA and HB) are effectively locked, leading to gel-like properties (**Figure 5a, Figure S5**). Once this strain is released, the material returns to the equilibrium fluid state (**Figure 5b**, +10 min deformation).

The dynamic stress response and genetic encodability of the noncovalent network formed by C2 and C5 through the LHD101 heterodimer make such materials attractive for applications in synthetic biology. Recent studies have used pairs of naturally occurring proteins to make intracellular hydrogels that mimic the function of RNA granules (45). We hypothesized that the analogous noncovalent interaction between the constituents (*i.e*., C2-5-HA and C5-5-HB) would lead to the intracellular formation of complex coacervates (non-Newtonian droplets). To explore the behavior of the material in living mammalian cells, we co-expressed C2-5-HA and C5-5-HB-GFP in HeLa cells, where green fluorescent protein (GFP) was genetically fused to the latter to monitor assembly formation by fluorescent microscopy. When both network constituents were coexpressed, we observed intracellular droplet-like punctae, demonstrating intracellular networking and complex coacervation formation through noncovalent interactions (**Figure 5d**). In contrast, the expression of C5-5-HB-GFP alone generated diffuse cytoplasmic fluorescence. Moreover, we found that intracellular droplet formation required a certain threshold concentration of both components. We co-expressed C2-5-HA-mCherry and C5-5-HB-GFP in HEK293T cells to determine the phase transition of the components from freely diffused proteins to self-assembled droplet punctae. As shown in **Figure S6**, droplet formation required a certain intensity of both C2-5-HA-mCherry and C5-5-HB-GFP. To assess the fluid properties of the droplet, we performed fluorescence recovery after photobleaching (FRAP) experiments, wherein a part of a droplet was photobleached and then allowed to recover fluorescence over time. The fluorescence recovery half-life time of the C2-5-HA × C5-5-HB-GFP droplet was 7.0 sec (**Figure 5e-f, Supplementary Movie 1**) indicating that the assemblies are not rigid hydrogels but are fluids, with their constituents reorganizing over time, similar to our rheological results. In comparison, photobleaching of control cells singly transfected with C5-5-HB-GFP showed a faster recovery half-life time of ∼4.1 sec, presumably due to free diffusion of the C5 assemblies when not crosslinked into a network (**Figure 5f, Figure S7**, and **Supplementary Movie 2**). The fluorescence intensity loss after recovery of the C2-5-HA × C5-5-HB-GFP droplet matched the fraction of the droplet remaining after bleaching, indicating that the reorganization of the C2-5-HA × C5-5-HB-GFP building blocks occurred within the droplet rather than exchange with the surrounding cytoplasm (**Figure S8**). In contrast, similar co-expression of covalently hydrogel-forming components in mammalian cells resulted in stable intracellular gels. The expression of C5-5-ST-GFP alone generated diffuse cytoplasmic fluorescence (recovery half-life time ∼2.17 sec) (**Figure 5f-g,** and **Supplementary Movie 4**), whereas the co-expression of C2-5-SC × C5-5-ST-GFP resulted in an intracellular hydrogel that did not recover (or recovered extremely slowly) fluorescence after photobleaching (**Figure 5f-g,** and **Supplementary Movie 3**), which may imply the stiffness of the gel.

## Designed intracellular *de novo* protein hydrogel mechanics correlates with extracellular gels

The C5-ST building blocks containing rigid arms formed covalent hydrogels with C2-SC, whose stiffness varied significantly with the change in rigid arm length (**Figure 2e-f**). Taking advantage of the distinct rheological properties offered by this series of hydrogel-forming building blocks, we co-expressed the constituents in mammalian cells to determine whether such hydrogels can be formed intracellularly and if their mechanical properties matched with the *ex cellulo* counterparts. Co-expression of C2-1-SC-mCherry with C5_2H_-1-ST-GFP in HEK293T cells resulted in diffuse fluorescence, whereas both C2-1-SC-mCherry × C5_7H_-1-ST-GFP and C2-1-SC-mCherry × C5_15H_-1-ST-GFP pairs showed droplet-like gel formation (**Figure 6a**). As the formation of hydrogel droplets was contingent on the strong expression of the hydrogel components at appropriate stoichiometries, hydrogels did not form in all cells expressing either the C2-mCherry or C5-GFP constructs. Therefore, we measured the propensity of intracellular gel formation, defined as the percentage of cells expressing either GFP or mCherry where the fluorescent proteins had formed intracellular condensates. The propensity of gel formation increased in the order of 2H<7H<15H (rigid arm length) (**Figure 6b**), exhibiting a strong positive correlation with rheologically measured bulk gel stiffnesses (**Figure 2e**). FRAP studies of these droplet gels further reflected their mechanical strength: 15H gels did not recover their fluorescence (**Supplementary Movie 7**) (recovery half-life time ∼6053 sec), whereas 7H gels recovered up to about 30% of their original intensity (**Supplementary Movie 6**) (recovery half-life time ∼3.23 sec). On the other hand, 2H assemblies failed to form any detectable gels and therefore recovered their fluorescence quickly (**Supplementary Movie 5**) (recovery half-life time ∼1.38 sec). Collectively, these experiments indicate that the 15H system may have formed gels with no dynamic reorganization and exchange of the constituents, 7H formed gels with reorganization of the constituents to some extent, whereas 2H failed to form any gels and therefore resulted in free flowing of the constituents (**Figure 6c, Figure S9**). This behavior of intracellular gel mechanics mirrored trends in *ex cellulo*-formed gel mechanics (**Figure 2e-f**).

**Figure 6:**
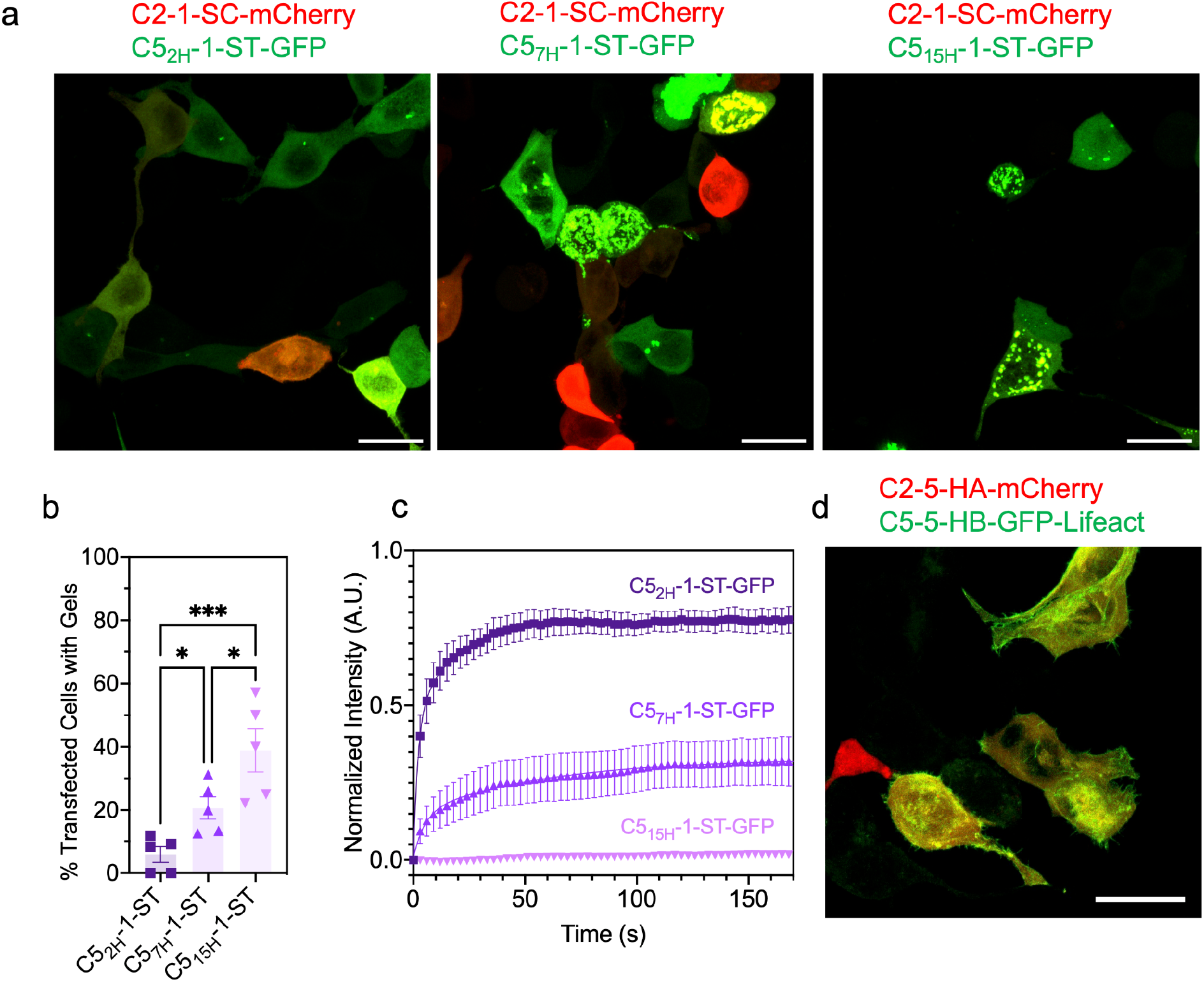
Intracellular gel mechanics correlates with properties of the corresponding extracellular gels. a) Intracellular gel formation between C2-1-SC-mCherry and C5-ST-GFP with varied rigid arm length (2H, 7H, and 15H). C5_7H_-1-ST-GFP and C5_15H_-1-ST-GFP formed efficient gels with C2-1-SC-mCherry, whereas C5_2H_-1-ST-GFP failed to do so. The propensity of intracellular gel formation of these combinations was quantified (b), which correlated with the stiffness of the corresponding extracellular gels (Figure 2e-f). c) FRAP experiments showing the recovery of fluorescence after photobleaching the intracellular gels. d) Targeting of intracellular gels to actin filaments through the fusion of a gel component to actin targeting Lifeact sequence (C5-5-HB-GFP-Lifeact). Scale bar 20 µm.

To further establish the functional relevance of these intracellular gels, we investigated whether protein assemblies could be localized within cells through genetic appendage of binding domains. We designed a C5 construct fused with the actin-binding Lifeact peptide (46). The resultant construct (*i.e*., C5-5-HB-GFP-Lifeact) was co-expressed with C2-5-HA-mCherry in HEK293T cells. Fluorescence microscopy imaging showed the colocalization of the C2-5-HA-mCherry (red) with C5-5-HB-GFP-Lifeact (green) throughout the cytoplasmic actin filaments with visible punctate-like structures indicating these gels may be targeted to the cytoskeleton (**Figure 6d**).

Intracellular liquid-liquid phase separated condensates have been generated from multivalent natural proteins with intrinsically disordered regions (47–50). Our de novo protein approach produces gels that resemble these condensates but goes beyond previous work by enabling finely tunable control over the fluid properties by varying the network chemistry and the structure of the components. This ability to finely modulate fluid properties could be useful for synthetic biology applications such as controlling enzyme sequestration, cell motility, and growth.

## Discussion

We have designed hydrogels from synthetic *de novo* protein building blocks with well-defined structures, valencies, and geometries connected in extended networks through either covalent or noncovalent interactions. Since the building blocks and interactions are well understood, our results provide a unique opportunity to relate the emergent macroscopic properties of hydrogels to the molecular and microscopic properties of the components. Full calculation of these properties would require extensive simulation efforts beyond the scope of this paper; here instead we have characterized the qualitative effects of varying building block topologies and association dynamics on hydrogel rheological properties. The dynamics of the component-component interaction driving hydrogel formation had a very large effect on the rheological properties. The covalent hydrogels formed using the SC/ST chemistry exhibited varied elastic moduli in a linker length between the core and the interaction domain and valency-dependent manner. In contrast, the non-covalent hydrogels formed using the reversibly associating designed LHD101 interaction exhibited dynamic viscoelastic properties in response to stress stimulus. Finally, simple changes to the secondary structure of gel constituents or network chemistry between them resulted in tunable biomaterials ranging in phase from fluid to gel, in storage moduli from soft to stiff, and generated both intra- and extracellularly.

There has been a growing interest in engineering extracellular matrices with defined stiffness for cellular differentiation and maintenance, and intracellular synthetic membrane-less organelles for a wide variety of cell functions. Our ability to fine-tune the elastic properties of protein networks by modulating the length of both flexible and rigid linkers, the connection valency, and the dynamics of the interactions mediating hydrogel crosslinking, while keeping a core structured region unaltered opens up avenues in synthetic biology and tissue engineering. The ability to genetically encode these networks extends the design space to within as well as between cells. Non-Newtonian shear-thickening has been observed in synthetic polymer-based materials (*i.e*., silly putty) and in certain biomolecules (*i.e.*, corn starch in water), however, to our knowledge, it is extremely rare in protein-based materials. More generally, assembling hydrogels from custom-designed components provides a systematic approach to relate microscopic structure to macroscopic properties, and as these connections become better understood, to custom design materials with desired viscoelasticity that can be formed both intra- and extracellularly.

## Methods

### Gene preparation

The amino acid sequences corresponding to respective oligomers (*i.e.,* C2, C5, T33, I53, LHD101, and SpyCatcher/Tag) were derived from previous reports and placed into pET29b+ vector with necessary modification, and incorporation of (GGS)_n_ linker, etc. For the full list of sequences, please see the supplementary information. For two-component designs (T33 and I53), all designs were expressed bi-cistronically by appending an additional ribosome binding site (RBS) in front of the second sequence, with only one of the components containing a 6xHis tag. Genes were synthesized by Integrated DNA Technologies (IDT).

### Protein expression and purification

All genes in the pET29b+ vector were transformed into *E. coli* cells [BL21 Lemo21 (DE3)] for expression. Proteins were expressed using an auto-induction protocol at 37 °C for 16–24 h on a 500 mL scale. Cells were harvested by centrifugation at 4000 × *g* for 10 min. Cell pellets were then resuspended in 25–30 mL lysis buffer (TBS, 25 mM Tris, 300 mM NaCl, pH8.0, 10 mM imidazole, 0.20 mg/mL DNase I) and sonicated for 2.5 min total on time at 80-90% power (10 s on/off) (QSonica). Lysates were then centrifuged at 14,000 × *g* for 30 min. Clarified lysates were then passed through Ni-NTA resin (QIAgen), washed with wash buffer (TBS, 25 mM Tris, 300 mM NaCl, pH 8.0, 40 mM imidazole), then eluted with elution buffer (TBS, 25 mM Tris, 300 mM NaCl, pH 8.0, 500 mM imidazole). Note that each protein was expressed in 12 × 500 mL cultures, and the eluents were collected together to concentrate with a 10,000 m/w cutoff spin concentrator (Millipore) to make the final concentration at least 100 mg/mL.

### Hydrogel preparation

Covalent hydrogels were formed by mixing SpyCatcher (SC)-modified C2 with SpyTagged (ST) higher-order oligomer in a 1:1 molar ratio of SC:ST, keeping the total w/v concentration fixed at 10%. Most constructs formed hydrogel immediately after mixing, but the samples were kept for ∼12 hrs for complete network formation. For visual inspection and gel photography, the mixtures were made in a cloning cylinder (4 mm inner diameter) in 40 microliter volume. The cylinder top was wrapped with a parafilm and kept at room temperature for 12 hours, and digital photographs were taken after carefully removing the cylinder. Note that, for the C2-(GGS)_n_-SC protein series, we only have C2-1-SC and C2-5-SC, whereas, for the C5-(GGS)_n_-ST series, we have n = 1 (C5-1-ST), 5 (C5-5-ST), and 10 (C5-10-ST). We did not include a C2-10-SC analog for this study as its solubility was poor and it did not reach the required 100 mg/mL concentration mark. Noncovalent hydrogels were made similar to the covalent at 20% w/v. For the deformation studies, C2-5-HA was mixed with C5-5-HB at an equimolar ratio of HA and HB on a parafilm sheet, and allowed to fully crosslink for half an hour. Pictures were taken 10 minutes before deformation, instantly after deformation, and 10 minutes after deformation.

### Rheometry

In situ rheology was conducted on an Anton Paar MCR 302 rheometer fitted with an 8mm parallel plate measurement attachment. Hydrogel droplets were formed at a volume of 30µL and thickness of 0.5mm on glass coverslips overnight in a humidified chamber prior to frequency sweep experiments. Gels on coverslips were then secured to the rheometer base plate for mechanical analysis. Frequency sweeps were conducted at a strain amplitude of 10% across frequencies from 0.1 to 500 rad/s unless otherwise stated. Linear viscoelasticity was verified in the strain amplitude domain by amplitude sweep rheology at frequencies in the linear viscoelastic region (1 rad/s for covalently crosslinked gels and 10 rad/s for noncovalently crosslinked gels) and amplitudes ranging from 1 to 100% strain. Cyclic strain time sweep experiments were conducted across 5 cycles of high strain (500% amplitude, 300 seconds) followed by a low-strain recovery period (1% amplitude, 1800 seconds).

### Cell encapsulation

10T1/2 cells were mixed with gel precursors to form a 10% w/v (in 1X PBS) hydrogel containing 10^6^ cells/mL, of which 10µL droplets were allowed to polymerize at 37°C in a humidified 96-well tissue culture dish. After 60 minutes of polymerization, media was added. At the specified timepoints, gels were stained with a LIVE/DEAD Cell Imaging kit (Thermo Fisher) per the manufacturer’s instructions and imaged immediately on a Leica Stellaris 5 confocal microscope under 10x magnification. Viable and nonviable cells were counted using CellProfiler 4.0.

### Intracellular assembly

All genes (*i.e*., C5-5-HB-GFP, C5-5-HB-GFP-Lifeact, C2-5-HA, C2-5-HA-mCherry, C5-5-ST-GFP, C2-5-SC, C2-1-SC-mCherry, C5_2H_-1-ST-GFP, C5_7H_-1-ST-GFP, and C5_15H_-1-ST-GFP) were cloned into the pcDNA3.1+ mammalian expression vector, which were transfected into HeLa cells (ATCC CCL-2) or HEK293T [cultured in Dulbecco’s modified Eagle’s medium (DMEM) at 37°C and 5% CO_2_] in the right combination. Briefly, cells were plated at 20,000 cells per well in Cellview cell culture slides (Greiner Bio-One ref 543079), then twenty-four hours later, cells were transiently transfected at a concentration of 140 ng total DNA per well and 1 μg/μl PEI-MAX (Polyscience) mixed with Opti-MEM medium (Gibco). Transfected cells were incubated at 37°C and 5% CO2 for 24 to 36 hours before being imaged.

### FRAP imaging

Bleaching movies were acquired with a commercial OMX-SR system (GE Healthcare) equipped with a Toptica diode 488 nm and 405 nm lasers and stage/objective heating set to 37 degrees C. Emission was collected on a PCO.edge sCMOS camera using an Olympus 60X 1.49NA ApoN oil immersion TIRF lens. 1024×1024 images (pixel size 6.5 μm) were captured with no binning. The acquisition was controlled with AcquireSR Acquisition control software. Photobleaching was carried out by acquiring a reference image followed by bleaching of a spot using a 600 ms pulse from a 405 nm laser. Recovery was monitored by imaging every 3 (rigid arm series) or 15 seconds for 3 minutes.

For FRAP analysis, FRAP data were plotted and fit to a single exponential association model using GraphPad Prism Version 9.3.1. Three ROIs of the same size were drawn on each image to measure (a) FRAP spot intensity (I_FRAP_) (b) a reference spot for photobleaching correction (I_Ref_), and (c) a baseline spot outside the cell for background correction (I_Base_). The baseline intensity was first subtracted from both I_FRAP_ and I_Ref_ to correct for image background(51).

I_FRAP Corr_ = I_FRAP_ –

I_Base_ I_Ref Corr_ = I_Ref_ - I_Base_

Next, correction for bleaching and normalization of intensities was performed using the following equation:

I_Norm_ = (I_Ref_pre_/I_Ref_corr(t)_) * (I_FRAP_corr(t)_/I_FRAP_pre_)

Where I_Ref_pre_ and I_FRAP_pre_ are the pre-bleaching background-corrected intensities and I_Ref_corr(t)_ and I_FRAP_corr(t)_ are the background-corrected intensities at each time point.

Finally, all traces were averaged and fitted to exponential association equations giving the half-time of fluorescence recovery and immobile fraction.

### Molecular dynamics

Simulations were conducted using molecular dynamics (MD) implemented in the HOOMD-blue particle simulation package (https://github.com/glotzerlab/hoomd-blue) (39). In the simulation model, C5 is represented as a rigid body of spheres with five-fold symmetry, which is connected by a flexible GGS linkers and SpyTag (**Figure S2 and Table S1**). The beads of the flexible linkers are bonded each other by a harmonic potential, 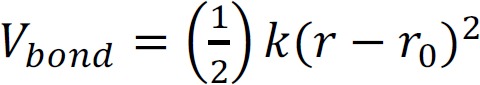, where *k* = 300ε/σ^2^ and *r*_0_ = 1.0σ. Here, *r* is the distance between the centers of beads of the GGS linker, σ is the distance unit, and ε is the energy unit in simulation. Similarly, C2 is represented as a rigid body of spheres that are linearly connected, and the rigid body of C2 is connected by flexible GGS linkers and SpyCatcher body (**Figure S2b**). The SpyCatcher body is represented as a rigid body of spheres that are connected in a cylindrical shape, and there are SpyCatcher patches at the center of the cylinder (**Figure S2b, c**), and the patches interact with SpyTag of C5 through an attractive Gaussian potential, 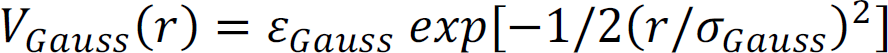, where 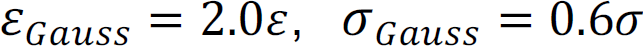, and *r* is the distance between the center of a bead of SpyTag and the center of a SpyCatcher.patch. All beads interact with each other (except SpyTag – SpyCatch patch pairs) via a purely repulsive WCA potential (52) to avoid overlap:

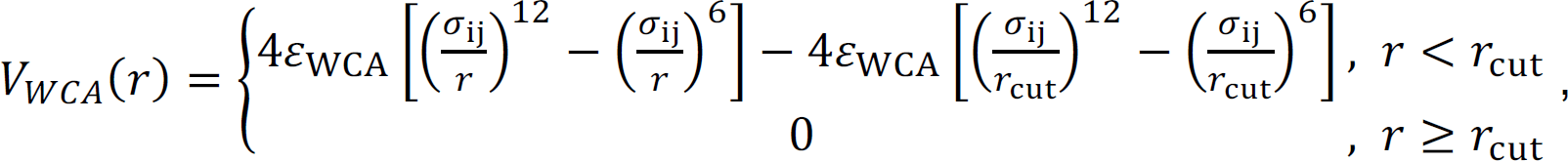

where ε_*WCA*_ = 0.1ε, σ_*ij*_ = *r_i_* + *r_j_*, *r*_*cut*_ = σ_*ij*_ × 2^1/6^, *i* and *j* are the type of beads that are interacting with, and *r_j_* is the radius of *i* bead listed in Table S1. We initialized system of *N*_*C*5_ = 400 and *N*_*C*2_ = 1,000 in a periodic box with a very dilute condition, *ϕ* = (*V*_*C*5_ + *V*_*C*2_)/*V*_*Box*_∼6.2 × 10^−4^, where *V*_*C*5_, *V*_*C*2_, *V*_>“?_ are the volume of C5, C2 and simulation box, respectively. The *V*_*C*5_ and *V*_*C*2_ do not include the volume of the GGS linker to have consistency throughout all systems. The initial positions of C5 and C2 are randomly assigned within the simulation box without making overlap. Various C2-*n* × C5-*m* combinations were used for each system, where *n* and *m* are the length of the GGS linker and *n* = 1,3,5,10 and *m* = 1,3,5,10. We compressed the system to *ϕ* = 0.023 for 10^=^ MD timesteps and thermalized for 5 × 10^B^ MD timesteps in NVT ensemble (*T*^∗^ = 0.2 *kT*/ε), where *T*^∗^ is the reduced temperature and each MD timestep is 0.002 for all systems. The density distribution of the simulated hydrogels (**Figure 4b**) was computed using GaussianDensity module implemented in freud python package (53). The coordinates of every C5 body, C2 body and SpyCatcher body were used for the density calculation. The g(r) (**Figure 4c**) of the system was computed based on the center of mass of C5 and C2.

## ACKNOWLEDGMENTS

This work was supported by the National Science Foundation (NSF) award CHE-1629214 (D.B.), a generous gift from the Audacious Project (D.B., N.I.E., S.L.), the Open Philanthropy Project (D.B., Y.H., R.M.), the Wu Tsai Translational Investigator Fund at the Institute for Protein Design (G.U.), the US DOE BES Energy Frontier Research Center CSSAS (The Center for the Science of Synthesis Across Scales) located at the University of Washington award number DESC0019288 (D.B., R.M.). R.M. and D.D.S. are recipients of Washington Research Foundation (WRF) Innovation fellowships. R.C.B acknowledges funding from the NSF Graduate Research Fellowship Program (DGE 176211) and UW ISCRM fellows program. This work was also supported through a Faculty Early Career Development (CAREER) award DMR 1652141 (C.A.D.) from the NSF, as well as a Maximizing Investigators’ Research Award (R35GM138036 to C.A.D.) from the National Institutes of Health (NIH). R.S. acknowledges NSF awards 2036803 and 1830893. R.M. acknowledges S. Cem Millik (University of Washington) for valuable discussions.

## Author contribution

R.M., Y.H., R.S., and D.B. conceived the idea. R.M., R.C.B., C.A.D., and D.B. designed experiments and wrote the manuscript with contributions from other authors. R.M. designed, purified, and characterized proteins; made hydrogels for imaging and rheometry; and analyzed data. R.C.B. performed rheometry, cell encapsulations and analyzed data. N.I.E. helped in protein production. S.L. performed molecular dynamics simulations and contributed to the manuscript writing. J.D. performed microscopy experiments, FRAP analysis. M.A. performed mammalian cell culture and transfection experiments. D.D.S. assisted in the heterodimer generation. G.U. provided suggestions for protein building blocks. N.G. and A.S. expressed and purified proteins and DNA constructs. R.S., C.A.D., and D.B. supervised the project.

