## Supplementary information for "De novo design of modular protein hydrogels with programmable intra- and extracellular viscoelasticity"

[1]Department of Biochemistry, University of Washington, Seattle, WA 98195
[2]Institute for Protein Design, University of Washington, Seattle, WA 98195
[3]Harvard Medical School, and Stem Cell Program at Boston Children's Hospital, Boston, MA 02115
[4]Department of Bioengineering, University of Washington, Seattle, WA 98195
[5]Department of Chemical Engineering, University of Washington, Seattle, WA 98195
[6]Institute for Stem Cell and Regenerative Medicine, University of Washington, Seattle, WA 98195
[7]Department of Chemistry, University of Washington, Seattle, WA 98195
[8]Molecular Engineering & Sciences Institute, University of Washington, Seattle, WA 98195
[9]Molecular and Cellular Biology Graduate Program, University of Washington, Seattle, WA 98195
[10]Medical Scientist Training Program, University of Washington, Seattle, WA 98195
[11]Howard Hughes Medical Institute, University of Washington, Seattle, WA 98195
[12]Department of Chemical and Biomolecular Engineering, Johns Hopkins University, Baltimore, MD 21218
[13]Department of Computer Science, Johns Hopkins University, Baltimore, MD 21218
[#]These authors contributed equally: Rubul Mout, Ross Bretherton

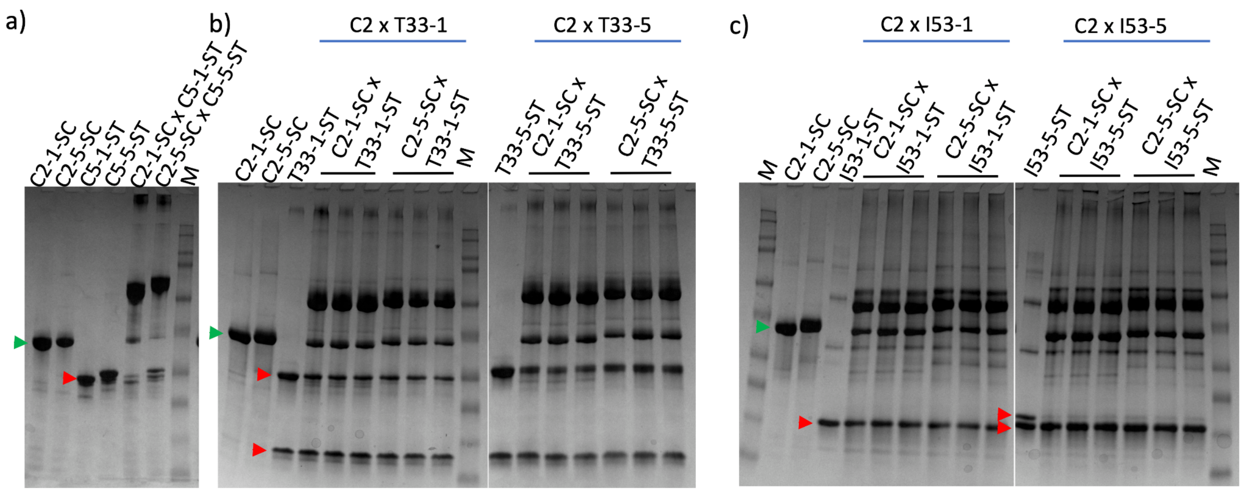


**Figure S1**. SDS-PAGE gel showing covalent crosslinking between SpyCatcher-modified C2 (blue arrow) and SpyTagged higher-order oligomers (red arrow). Upon covalent network formation through SpyLigation, the conjugated oligomers shift in their position on the gel. a) C2 x C5; b) C2 x T33; and c) C2 x I53 hydrogels. Note that T33 and I53 oligomers are constituted of two components and hence two bands are observed in the gel. Moreover, in I53-1-ST, both the components have identical sizes and the two bands are not distinguishable. T33 and I53 samples were run in triplicate (b, c). M: standard protein ladder.

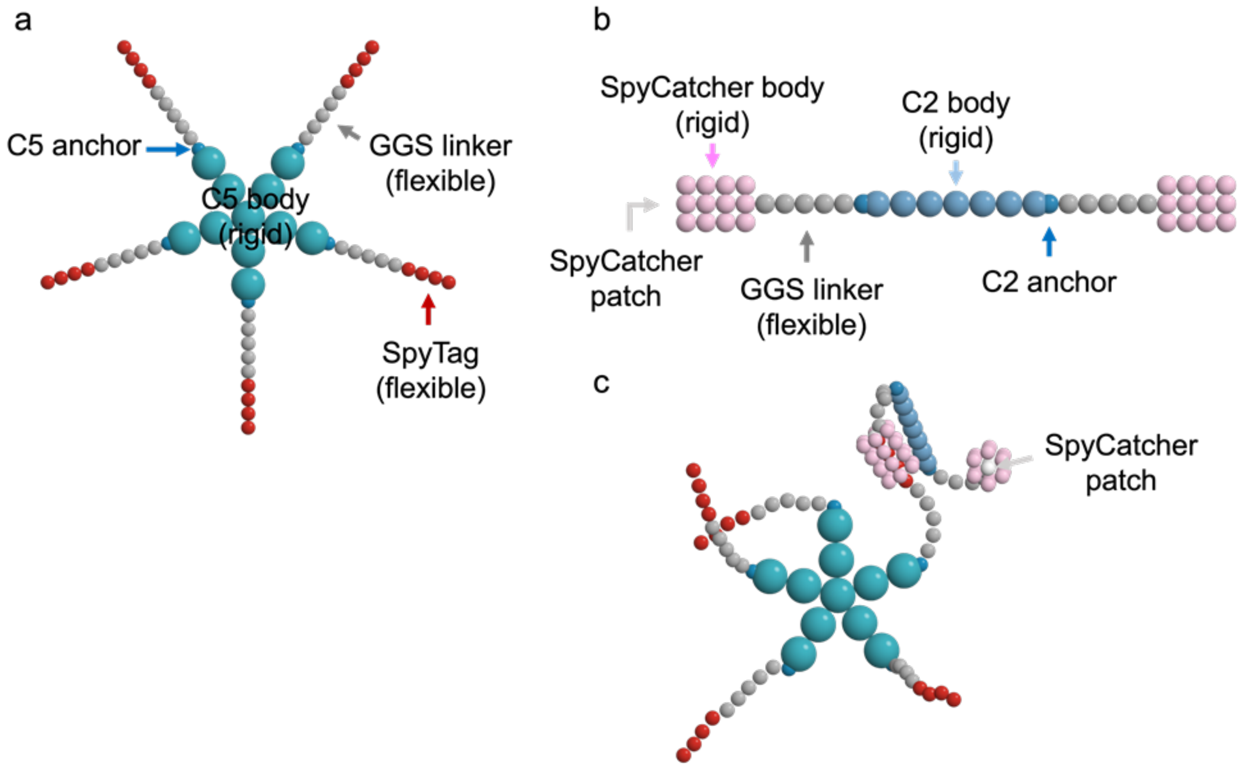


**Figure S2.** MD simulation model with representative examples C5-5-ST and C2-5-SC. (**a**) C5 consists of C5 body, C5 anchor, GGS linker, and SpyTag, and their coordinates are listed in Table

S1. (**b**) C2 consists of C2 body, C2 anchor, GGS linker, SpyCatcher body, and SpyCatcher patch. (**c**) An attractive potential is applied between SpyTag and SpyCatcher patch so that the SpyTag can go into the SpyCatcher body.

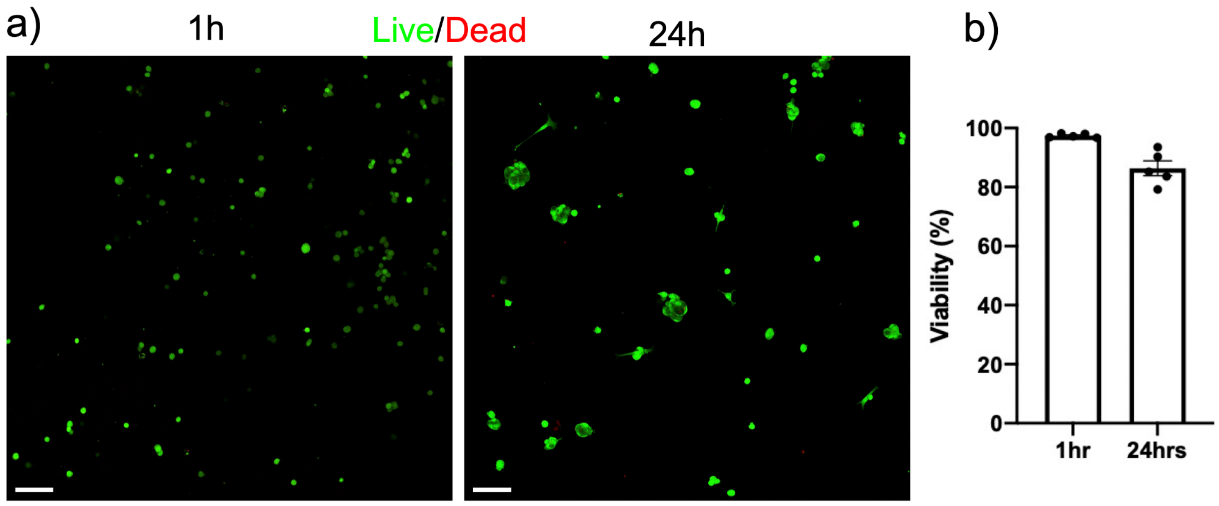


**Figure S3**: Cell encapsulation of 10T1/2 fibroblast cells in C2-5-SC x C5-5-ST gels. a) Cells were encapsulated and grown for 1 and 24h. Cell viability was assessed by LIVE/DEAD staining and confocal imaging. b) Percentage of cell viability was quantified by CellProfiler. Scale bar 20 μm.

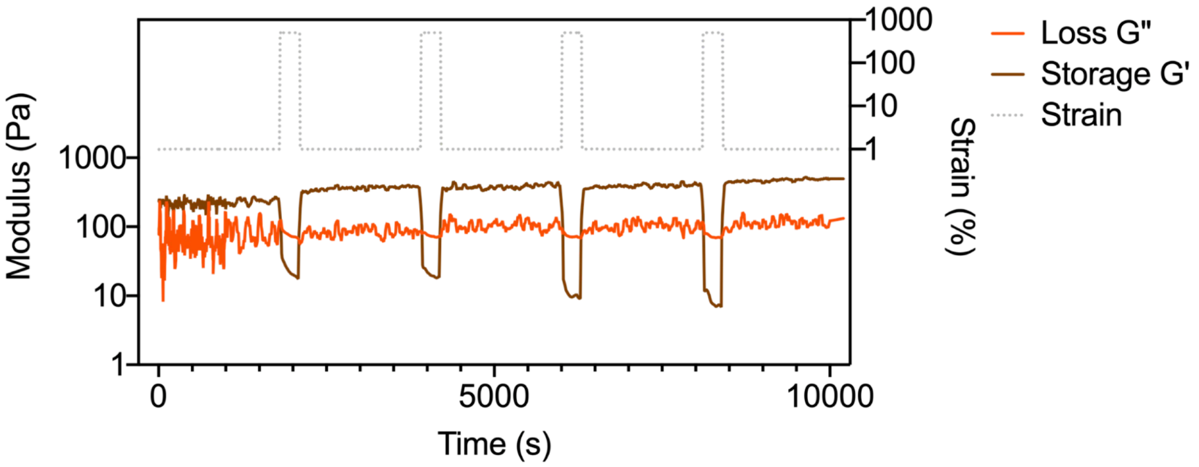

**Figure S4.** Self-healing properties of C2-5-HA x C5-5-HB noncovalent protein networks mediated by LHD101 heterodimer. The gel networks were capable of self-healing over multiple cycles of 500% strain.

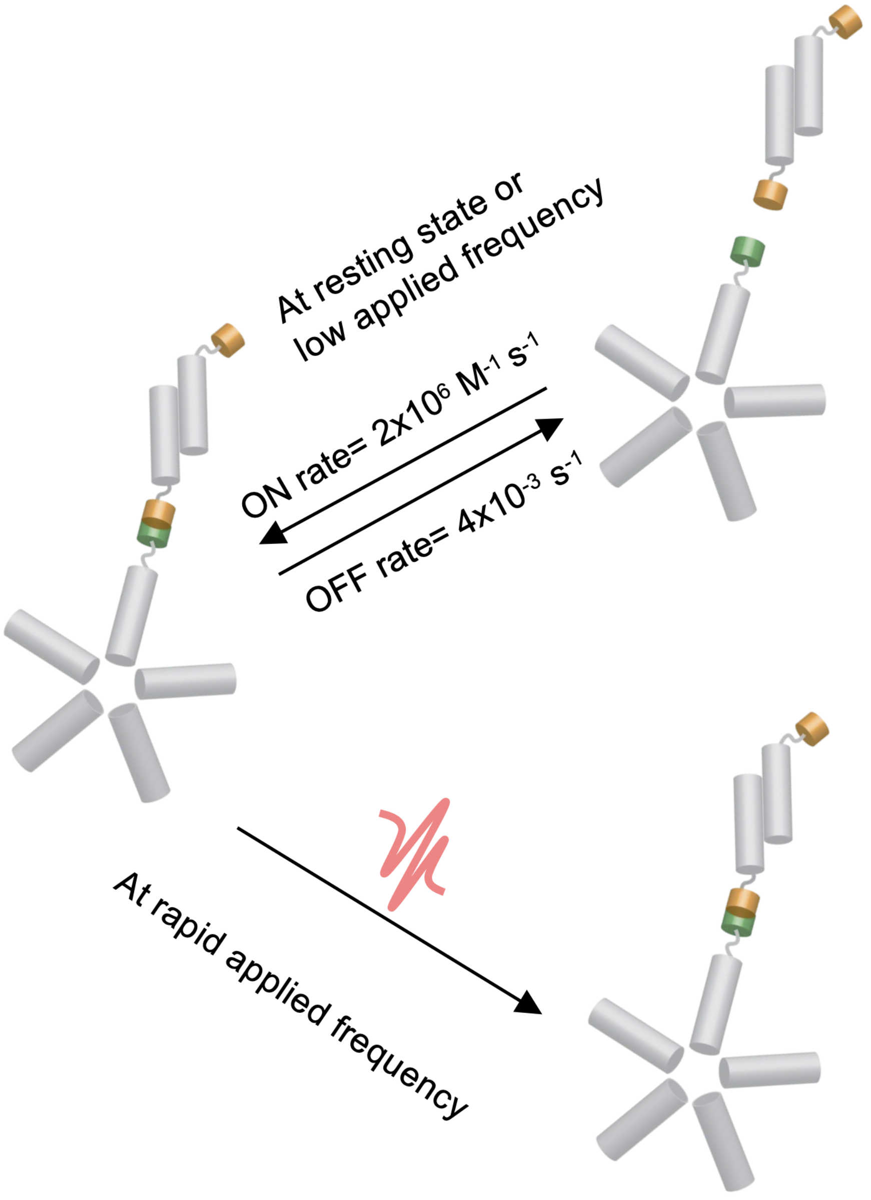


**Figure S5.** Model showing the noncovalent assembly between C2-5-HA and C5-5-HB at a resting state or at a state when the applied frequency is low vs when the applied frequency is rapid. At the resting state, due to the rapid ON/OFF rate of the heterodimer units, the material behaves like a liquid, whereas, if the assembly exerts a deformation frequency faster than the heterodimer LHD101A:B (HA and HB) ON/OFF rate then the intermolecular interactions get 'locked', and therefore behaves like a gel.

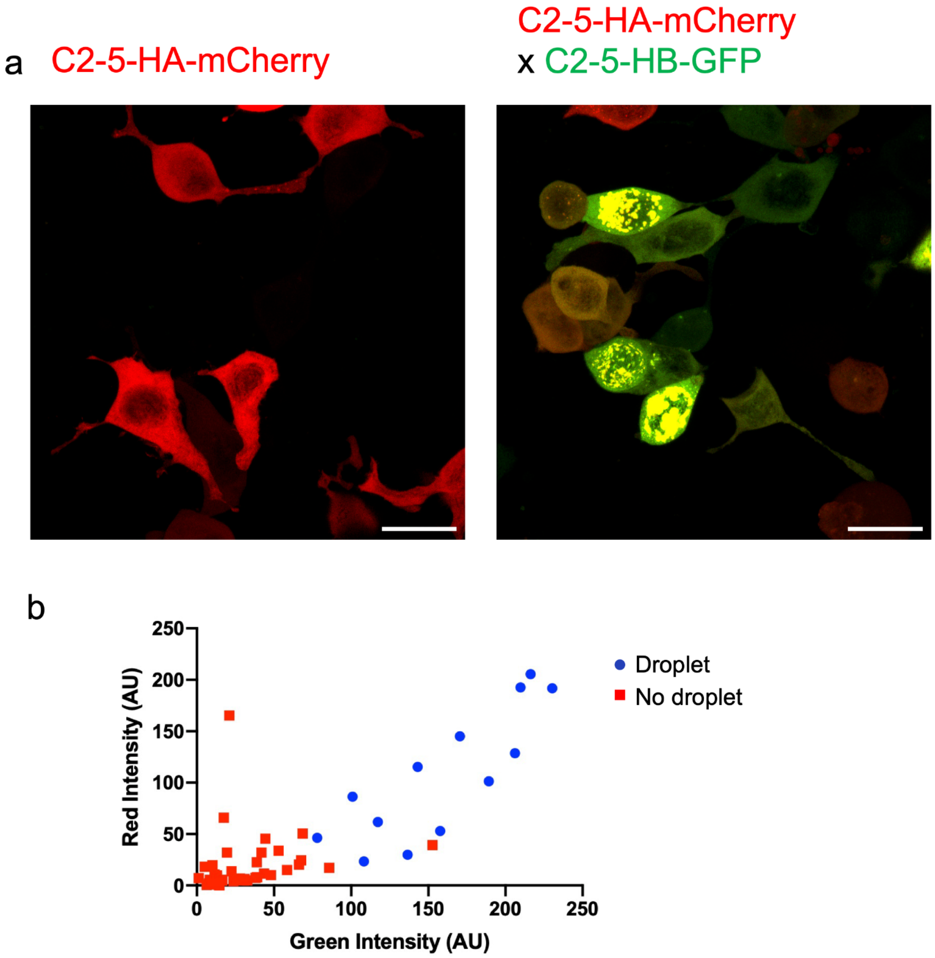


**Figure S6**. The phase transition from freely diffused constituents to droplet punctae. a) Expression of C2-5-HA-mCherry alone or the coexpression of C2-5-HA-mCherry and C2-5-HB-GFP. b) Droplet formation occurred only at a certain threshold expression of both the components, measured through red (mCherry) and green (GFP) fluorescent intensity.

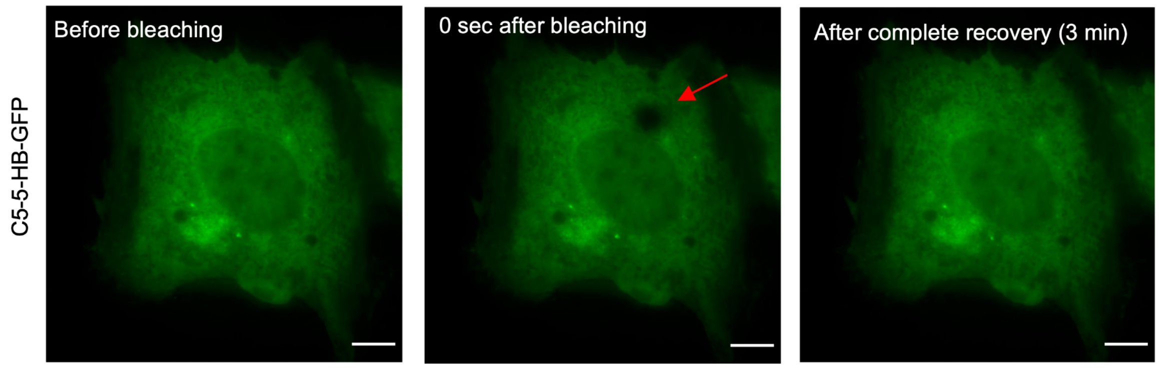


**Figure S7**. Fluorescence recovery after photobleaching (FRAP) experiment of the control C5-5-HB-GFP expressing cells demonstrated a rapid recovery in the absence of a droplet formation.

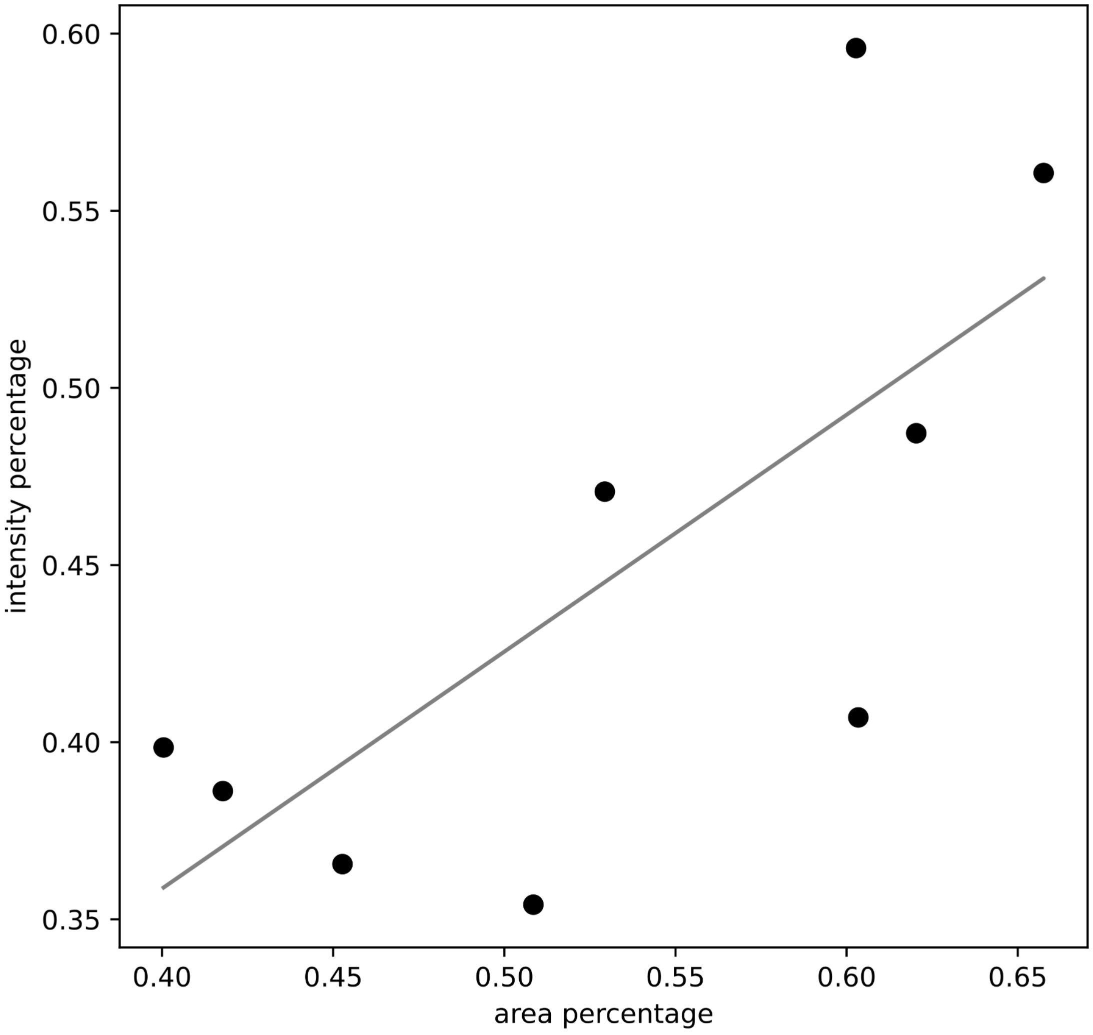

**Figure S8.** Measurement of bleached area versus fluorescence intensity recovery. The x-axis represents the area percentage, measured by taking the ratio of the droplet area pre and post bleach; whereas the y-axis represents the intensity percentage, measured by taking the ratio of full droplet intensity pre-bleach to the intensity post-recovery. For most parts, a linear correlation between the bleached area and the recovery intensity percentage was observed.

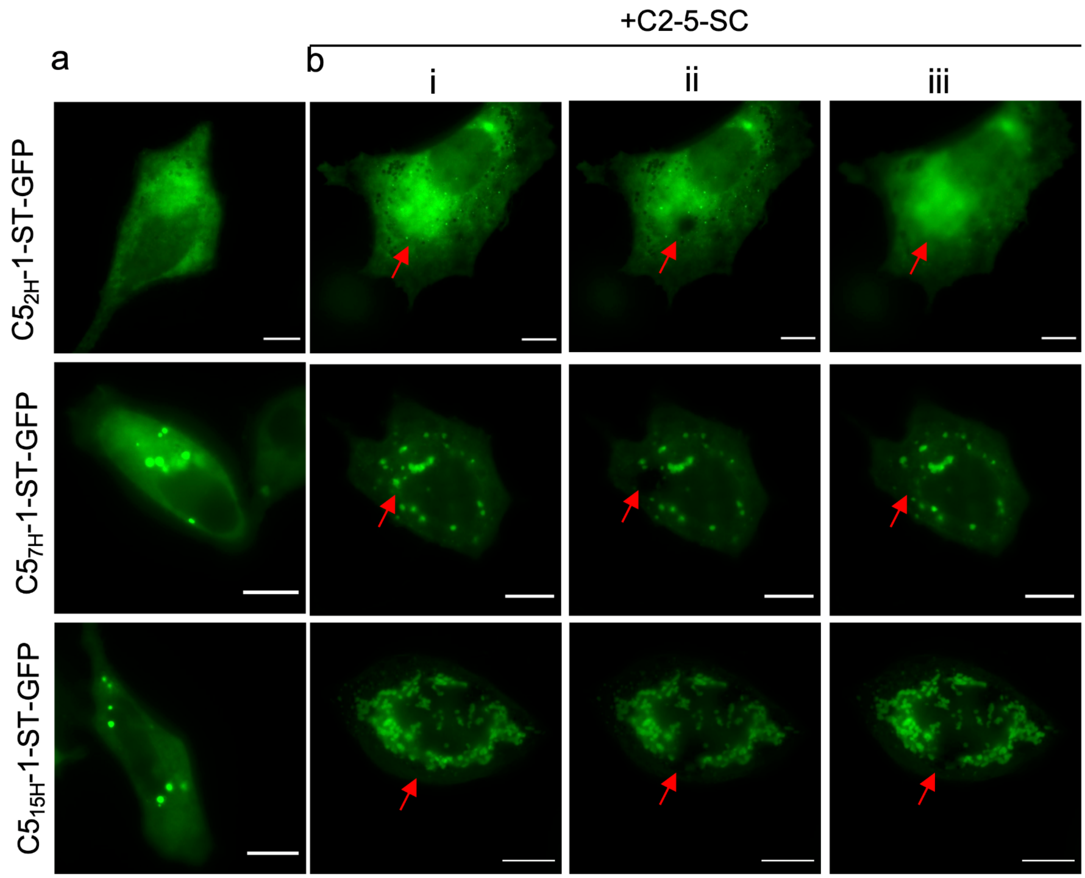


**Figure S9**: Fluorescence recovery after photobleaching (FRAP) experiment of the rigid arm series. a) Expression of C5$_{2H}$-1-ST-GFP, C5$_{7H}$-1-ST-GFP, and C5$_{15H}$-1-ST-GFP constructs alone, or with C2-5-SC (b). (i) Before photobleaching, (ii) right after photobleaching, and (iii) 3 minutes after photobleaching. Scale bar 10 μm.

**Table S1.** Coordinates (XYZ) of components comprising C5-5-ST and C2-5-SC models. The radius of each bead of the C5 body is 1.2$\sigma$, and that of the C2 body is 0.67$\sigma$. The radius of all other beads is 0.5$\sigma$. The length of the GGS linker (*i.e.* the number of beads of the GGS linker) can be changed depending on the system.

| C5-5 | | | | C2-5 | | | |
|---|---|---|---|---|---|---|---|
| Type | x | y | z | Type | x | y | z |
| C5 body | 0.000 | -0.002 | 0.000 | C2 anchor | -5.466 | 0.000 | -0.059 |
| C5 body | 0.000 | 2.390 | 0.000 | C2 anchor | 3.934 | 0.000 | -0.059 |
| C5 body | 0.000 | 4.782 | 0.000 | C2 body | -4.795 | 0.000 | -0.059 |
| C5 body | 2.275 | 0.738 | 0.000 | C2 body | -3.452 | 0.000 | -0.059 |
| C5 body | 4.550 | 1.477 | 0.000 | C2 body | -2.109 | 0.000 | -0.059 |
| C5 body | 1.406 | -1.937 | 0.000 | C2 body | -0.766 | 0.000 | -0.059 |
| C5 body | 2.812 | -3.872 | 0.000 | C2 body | 0.576 | 0.000 | -0.059 |
| C5 body | -1.406 | -1.937 | 0.000 | C2 body | 1.919 | 0.000 | -0.059 |
| C5 body | -2.812 | -3.872 | 0.000 | C2 body | 3.262 | 0.000 | -0.059 |
| C5 body | -2.275 | 0.738 | 0.000 | GGS linker | -6.466 | 0.000 | -0.059 |
| C5 body | -4.550 | 1.477 | 0.000 | GGS linker | -7.466 | 0.000 | -0.059 |
| C5 anchor | 0.000 | 5.978 | 0.000 | GGS linker | -8.466 | 0.000 | -0.059 |
| C5 anchor | 5.687 | 1.846 | 0.000 | GGS linker | -9.466 | 0.000 | -0.059 |
| C5 anchor | 3.515 | -4.840 | 0.000 | GGS linker | -10.466 | 0.000 | -0.059 |
| C5 anchor | -3.515 | -4.840 | 0.000 | GGS linker | 4.934 | 0.000 | -0.059 |
| C5 anchor | -5.687 | 1.846 | 0.000 | GGS linker | 5.934 | 0.000 | -0.059 |

| | | | | | | | |
|---|---|---|---|---|---|---|---|
| GGS linker | 0.000 | 6.978 | 0.000 | GGS linker | 6.934 | 0.000 | -0.059 |
| GGS linker | 0.000 | 7.978 | 0.000 | GGS linker | 7.934 | 0.000 | -0.059 |
| GGS linker | 0.000 | 8.978 | 0.000 | GGS linker | 8.934 | 0.000 | -0.059 |
| GGS linker | 0.000 | 9.978 | 0.000 | SpyCatcher patch | -11.466 | 0.000 | -0.059 |
| GGS linker | 0.000 | 10.978 | 0.000 | SpyCatcher patch | -12.466 | 0.000 | -0.059 |
| GGS linker | 6.638 | 2.155 | 0.000 | SpyCatcher patch | -13.466 | 0.000 | -0.059 |
| GGS linker | 7.589 | 2.464 | 0.000 | SpyCatcher patch | -14.466 | 0.000 | -0.059 |
| GGS linker | 8.540 | 2.773 | 0.000 | SpyCatcher patch | 12.934 | 0.000 | -0.059 |
| GGS linker | 9.492 | 3.082 | 0.000 | SpyCatcher patch | 11.934 | 0.000 | -0.059 |
| GGS linker | 10.443 | 3.391 | 0.000 | SpyCatcher patch | 10.934 | 0.000 | -0.059 |
| GGS linker | 4.103 | -5.649 | 0.000 | SpyCatcher patch | 9.934 | 0.000 | -0.059 |
| GGS linker | 4.691 | -6.458 | 0.000 | SpyCatcher body | -11.466 | 0.000 | 1.041 |
| GGS linker | 5.278 | -7.267 | 0.000 | SpyCatcher body | -11.466 | 0.953 | 0.491 |
| GGS linker | 5.866 | -8.076 | 0.000 | SpyCatcher body | -11.466 | 0.953 | -0.609 |
| GGS linker | 6.454 | -8.885 | 0.000 | SpyCatcher body | -11.466 | -0.953 | -0.609 |

| | | | | | | | |
|---|---|---|---|---|---|---|---|
| GGS linker | -4.103 | -5.649 | 0.000 | SpyCatcher body | -11.466 | -0.953 | 0.491 |
| GGS linker | -4.691 | -6.458 | 0.000 | SpyCatcher body | -12.466 | 0.000 | 1.041 |
| GGS linker | -5.278 | -7.267 | 0.000 | SpyCatcher body | -12.466 | 0.953 | 0.491 |
| GGS linker | -5.866 | -8.076 | 0.000 | SpyCatcher body | -12.466 | 0.953 | -0.609 |
| GGS linker | -6.454 | -8.885 | 0.000 | SpyCatcher body | -12.466 | -0.953 | -0.609 |
| GGS linker | -6.638 | 2.155 | 0.000 | SpyCatcher body | -12.466 | -0.953 | 0.491 |
| GGS linker | -7.589 | 2.464 | 0.000 | SpyCatcher body | -13.466 | 0.000 | 1.041 |
| GGS linker | -8.540 | 2.773 | 0.000 | SpyCatcher body | -13.466 | 0.953 | 0.491 |
| GGS linker | -9.492 | 3.082 | 0.000 | SpyCatcher body | -13.466 | 0.953 | -0.609 |
| GGS linker | -10.443 | 3.391 | 0.000 | SpyCatcher body | -13.466 | -0.953 | -0.609 |
| SpyTag | 0.000 | 11.978 | 0.000 | SpyCatcher body | -13.466 | -0.953 | 0.491 |
| SpyTag | 0.000 | 12.978 | 0.000 | SpyCatcher body | -14.466 | 0.000 | 1.041 |
| SpyTag | 0.000 | 13.978 | 0.000 | SpyCatcher body | -14.466 | 0.953 | 0.491 |
| SpyTag | 0.000 | 14.978 | 0.000 | SpyCatcher body | -14.466 | 0.953 | -0.609 |
| SpyTag | 11.394 | 3.700 | 0.000 | SpyCatcher body | -14.466 | -0.953 | -0.609 |
| SpyTag | 12.345 | 4.009 | 0.000 | SpyCatcher body | -14.466 | -0.953 | 0.491 |
| SpyTag | 13.296 | 4.318 | 0.000 | SpyCatcher body | 12.934 | 0.000 | 1.041 |

| | | | | | | | |
|---|---|---|---|---|---|---|---|
| SpyTag | 14.247 | 4.627 | 0.000 | SpyCatcher body | 12.934 | 0.953 | 0.491 |
| SpyTag | 7.042 | -9.694 | 0.000 | SpyCatcher body | 12.934 | 0.953 | -0.609 |
| SpyTag | 7.629 | -10.503 | 0.000 | SpyCatcher body | 12.934 | 0.000 | -1.159 |
| SpyTag | 8.217 | -11.312 | 0.000 | SpyCatcher body | 12.934 | -0.953 | -0.609 |
| SpyTag | 8.805 | -12.121 | 0.000 | SpyCatcher body | 12.934 | -0.953 | 0.491 |
| SpyTag | -7.042 | -9.694 | 0.000 | SpyCatcher body | 11.934 | 0.000 | 1.041 |
| SpyTag | -7.629 | -10.503 | 0.000 | SpyCatcher body | 11.934 | 0.953 | 0.491 |
| SpyTag | -8.217 | -11.312 | 0.000 | SpyCatcher body | 11.934 | 0.953 | -0.609 |
| SpyTag | -8.805 | -12.121 | 0.000 | SpyCatcher body | 11.934 | 0.000 | -1.159 |
| SpyTag | -11.394 | 3.700 | 0.000 | SpyCatcher body | 11.934 | -0.953 | -0.609 |
| SpyTag | -12.345 | 4.009 | 0.000 | SpyCatcher body | 11.934 | -0.953 | 0.491 |
| SpyTag | -13.296 | 4.318 | 0.000 | SpyCatcher body | 10.934 | 0.000 | 1.041 |
| SpyTag | -14.247 | 4.627 | 0.000 | SpyCatcher body | 10.934 | 0.953 | 0.491 |
| | | | | SpyCatcher body | 10.934 | 0.953 | -0.609 |
| | | | | SpyCatcher body | 10.934 | 0.000 | -1.159 |
| | | | | SpyCatcher body | 10.934 | -0.953 | -0.609 |
| | | | | SpyCatcher body | 10.934 | -0.953 | 0.491 |
| | | | | SpyCatcher body | 9.934 | 0.000 | 1.041 |
| | | | | SpyCatcher body | 9.934 | 0.953 | 0.491 |
| | | | | SpyCatcher body | 9.934 | 0.953 | -0.609 |
| | | | | SpyCatcher body | 9.934 | 0.000 | -1.159 |

| | | | | | | | |
|---|---|---|---|---|---|---|---|
| | | | | SpyCatcher body | 9.934 | -0.953 | -0.609 |
| | | | | SpyCatcher body | 9.934 | -0.953 | 0.491 |

**Amino acid sequence**: Yellow: linker; Red: SpyCatcher; Blue: SpyTag; Orange: LHD101A (HA); Purple: LHD101B (HB)

**Covalent**:

>C2-GGS-SpyCatcher (C2-1-SC)

**M**GTREEIIRELARSLAEQAELTARLERLLRELERLQREGSSDEDVRELLREIKELVREILKLIAEQILLI
AELLLAAIRSEAAELALRAIREAIELCKRSTDEELCQLLLRLALLLMELALLYPDSEAAKLALKAALEAI
ELCKQSTDEELCEELVKLAQKLIELAKRYPDSEAAKLALKAALEAIELCKQSTDEELCEELVKLAQKLIE
LAKRYPDSEEAKRALKEAKELIEQCKESTDEDECRELVKRAEELIREAKEGGSDSATHIKFSKRDIDGKE
LAGATMELRDSSGKTISTWISDGQVKDFYLMPGKYTFVETAAPDGYEIATAITFTVNEQGQVTVNGKATK
GGSWGLEHHHHHH

>C2-(GGS)$_5$-SpyCatcher (C2-5-SC)

**M**GTREEIIRELARSLAEQAELTARLERLLRELERLQREGSSDEDVRELLREIKELVREILKLIAEQILLI
AELLLAAIRSEAAELALRAIREAIELCKRSTDEELCQLLLRLALLLMELALLYPDSEAAKLALKAALEAI
ELCKQSTDEELCEELVKLAQKLIELAKRYPDSEAAKLALKAALEAIELCKQSTDEELCEELVKLAQKLIE
LAKRYPDSEEAKRALKEAKELIEQCKESTDEDECRELVKRAEELIREAKEGGSGGSGGSGGSGGSDSATH
IKFSKRDIDGKELAGATMELRDSSGKTISTWISDGQVKDFYLMPGKYTFVETAAPDGYEIATAITFTVNE
QGQVTVNGKATKGGSWGLEHHHHHH

>C5-(GGS)$_1$-SpyTag (C5-1-ST)

**M**GHHHHHHGWSGAHIVMVDAYKPTKGGSNDEKEKLKELLKRAEELAKSPDPEDLKEAVRLAEEVVRERPG
SNLAKKALEIILRAAEELAKLPDPEALKEAVKAAEKVVREQPGSNLAKKAQEIILRAAEELAKLEDEEAL
KEAIKAAEKVIELEPGSELAKEAKRIIEKAAKMLADILRKEMEKIREETEEVKKEIEESKKRPQSESAKN
LILIMQLLINQIRLLALQIRMLVLQLIL

>C5-(GGS)$_5$-SpyTag (C5-5-ST)

**M**GHHHHHHGWSGAHIVMVDAYKPTKGGSGGSGGSGGSGGSNDEKEKLKELLKRAEELAKSPDPEDLKEAV
RLAEEVVRERPGSNLAKKALEIILRAAEELAKLPDPEALKEAVKAAEKVVREQPGSNLAKKAQEIILRAA
EELAKLEDEEALKEAIKAAEKVIELEPGSELAKEAKRIIEKAAKMLADILRKEMEKIREETEEVKKEIEE
SKKRPQSESAKNLILIMQLLINQIRLLALQIRMLVLQLIL

>C5-(GGS)$_{10}$-SpyTag (C5-10-ST)

**M**GHHHHHHGWSGAHIVMVDAYKPTKGGSGGSGGSGGSGGSGGSGGSGGSGGSGGSNDEKEKLKELLKRAE
ELAKSPDPEDLKEAVRLAEEVVRERPGSNLAKKALEIILRAAEELAKLPDPEALKEAVKAAEKVVREQPG
SNLAKKAQEIILRAAEELAKLEDEEALKEAIKAAEKVIELEPGSELAKEAKRIIEKAAKMLADILRKEME
KIREETEEVKKEIEESKKRPQSESAKNLILIMQLLINQIRLLALQIRMLVLQLIL

>T33_chainA

**M**GEEAELAYLLGELAYKLGEYRIAIRAYRIALKRDPNNAEAWYNLGNAYYKQGDYDEAIEYYQKALELDP
NNAEAWYNLGNAYYKQGDYDEAIEYYEKALELDPENLEALQNLLNAMDKQG

>T33-(GGS)$_{1}$-SpyTag_chainB (T33-1-ST_chainB)

**M**IEEVVAEMIDILAESSKKSIEELARAADNKTTEKAVAEAIEEIARLATAAIQLIEALAKNLASEEFMAR
AISAIAELAKKAIEAIYRLADNHTTDTFMARAIAAIANLAVTAILAIAALASNHTTEEFMARAISAIAEL
AKKAIEAIYRLADNHTTDKFMAAAIEAIALLATLAILAIALLASNHTTEKFMARAIMAIAILAAKAIEAI
YRLADNHTSPTYIEKAIEAIEKIARKAIKAIEMLAKNITTEEYKEKAKKIIDIIRKLAKMAIKKLEDNRT
GGSAHIVMVDAYKPTKGSWGLEHHHHHH

>T33-(GGS)$_{5}$-SpyTag_chainB (T33-5-ST_chainB)

**M**IEEVVAEMIDILAESSKKSIEELARAADNKTTEKAVAEAIEEIARLATAAIQLIEALAKNLASEEFMAR
AISAIAELAKKAIEAIYRLADNHTTDTFMARAIAAIANLAVTAILAIAALASNHTTEEFMARAISAIAEL
AKKAIEAIYRLADNHTTDKFMAAAIEAIALLATLAILAIALLASNHTTEKFMARAIMAIAILAAKAIEAI
YRLADNHTSPTYIEKAIEAIEKIARKAIKAIEMLAKNITTEEYKEKAKKIIDIIRKLAKMAIKKLEDNRT
GGSGGSGGSGGSGGSAHIVMVDAYKPTKGSWGLEHHHHHH

>T33-(GGS)$_{10}$-SpyTag_chainB (T33-10-ST_chainB)

**M**IEEVVAEMIDILAESSKKSIEELARAADNKTTEKAVAEAIEEIARLATAAIQLIEALAKNLASEEFMAR
AISAIAELAKKAIEAIYRLADNHTTDTFMARAIAAIANLAVTAILAIAALASNHTTEEFMARAISAIAEL
AKKAIEAIYRLADNHTTDKFMAAAIEAIALLATLAILAIALLASNHTTEKFMARAIMAIAILAAKAIEAI
YRLADNHTSPTYIEKAIEAIEKIARKAIKAIEMLAKNITTEEYKEKAKKIIDIIRKLAKMAIKKLEDNRT
GGSGGSGGSGGSGGSGGSGGSGGSGGSGGSAHIVMVDAYKPTKGSWGLEHHHHHH

>I53_chainA

**M**GKYDGSKLRIGILHARWNAEIILALVLGALKRLQEFGVKRENIIIETVPGSFELPYGSKLFVEKQKRLG
KPLDAIIPIGVLIKGSTMHFEYICDSTTHQLMKLNFELGIPVIFGVLTCLTDEQAEARAGLIEGKMHNHG
EDWGAAAVEMATKFN

>I53-(GGS)$_{1}$-SpyTag_chainB (I53-1-ST_chainB)

**M**EEAELAYLLGELAYKLGEYRIAIRAYRIALKRDPNNAEAWYNLGNAYYKQGRYREAIEYYQKALELDPN
NAEAWYNLGNAYYERGEYEEAIEYYRKALRLDPNNADAMQNLLNAKMREEGGSAHIVMVDAYKPTKGSWG
LEHHHHHH

>I53-(GGS)$_5$-SpyTag_chainB (I53-5-ST_chainB)

**M**EEAELAYLLGELAYKLGEYRIAIRAYRIALKRDPNNAEAWYNLGNAYYKQGRYREAIEYYQKALELDPN
NAEAWYNLGNAYYERGEYEEAIEYYRKALRLDPNNADAMQNLLNAKMREEGGSGGSGGSGGSGGSAHIVM
VDAYKPTKGSWGLEHHHHHH

>I53-(GGS)$_{10}$-SpyTag_chainB (I53-10-ST_chainB)

**M**EEAELAYLLGELAYKLGEYRIAIRAYRIALKRDPNNAEAWYNLGNAYYKQGRYREAIEYYQKALELDPN
NAEAWYNLGNAYYERGEYEEAIEYYRKALRLDPNNADAMQNLLNAKMREEGGSGGSGGSGGSGGSGGSGG
SGGSGGSGGSAHIVMVDAYKPTKGSWGLEHHHHHH

>C5$_{2H}$-(GGS)$_1$-SpyTag (C5$_{2H}$-1-ST)

**M**GHHHHHHGWSGAHIVMVDAYKPTKGGSTRRKQEMKRLKKEMEKIREETEEVKKEIEESKKRPQSESAKN
LILIMQLLINQIRLLALQIRMLALQLQE

>C5$_{7H}$-(GGS)$_1$-SpyTag (C5$_{7H}$-1-ST)

**M**GHHHHHHGWSGAHIVMVDAYKPTK**GGS**SDLQEVADRIVEQLKREGRSPEEARKEARRLIEEIKQSAGGD
SELIEVAVRIVKFLEEAGMSPSEAAKVAVELIERIRRAAGGDSELIEKAVRIVRRLERRGLSPAEAAKIA
VAIIAAEVLSREAEKIREETEEVKKEIEESKKRPQSESAKNLILIMQLLINQIRLLALQIQMLRLQLEL

>C5$_{15H}$-(GGS)$_1$-SpyTag (C5$_{15H}$-1-ST)

**M**GHHHHHHGWSGAHIVMVDAYKPTKGGSSDLQEVADRIVEQLKREGRSPEEARKEARRLIEEIKQSAGGD
SELIEVAVRIVKELEEQGRSPSEAAKEAVELIERIRRAAGGDSELIEVAVRIVKELEEQGRSPSEAAKEA
VELIERIRRAAGGDSELIEVAVRIVKELEEQGRSPSEAAKEAVELIERIRRAAGGDSELIEVAVRIVKEL
EEQGRSPSEAAKEAVELIERIRRAAGGDSELIEVAVRIVKFLEEAGMSPSEAAKVAVELIERIRRAAGGD
SELIEKAVRIVRRLERRGLSPAEAAKIAVAIIAAEVLSREAEKIREETEEVKKEIEESKKRPQSESAKNL
ILIMQLLINQIRLLALQIQMLRLQLEL

**Noncovalent:**

>C2-(GGS)$_5$-LHD101A (C2-5-HA)

**M**GTREEIIRELARSLAEQAELTARLERLLRELERLQREGSSDEDVRELLREIKELVREILKLIAEQILLI
AELLLAAIRSEAAELALRAIREAIELCKRSTDEELCQLLLRLALLLMELALLYPDSEAAKLALKAALEAI
ELCKQSTDEELCEELVKLAQKLIELAKRYPDSEAAKLALKAALEAIELCKQSTDEELCEELVKLAQKLIE
LAKRYPDSEEAKRALKEAKELIEQCKESTDEDECRELVKRAEELIREAKEGGSGGSGGSGGSGGSGRQEK

VLKSIEETVRKMGVTMETHRSGNEVKVVIKGLHIKQQRQLYRDVRETSKKQGVETEIEVEGDTVTIVVRE
GSWGLEHHHHHH

>C5-$(GGS)_5$-LHD101B (C5-5-HB)

**M**GHHHHHHGWSGGRQEKVLKSIEETVRKMGVTMETHRSGNEVKVVIKGLHESQQEQLLEDVLRTAEKQGV
RVRIRFKGDTVTIVVREGGSGGSGGSGGSGGSNDEKEKLKELLKRAEELAKSPDPEDLKEAVRLAEEVVR
ERPGSNLAKKALEIILRAAEELAKLPDPEALKEAVKAAEKVVREQPGSNLAKKAQEIILRAAEELAKLED
EEALKEAIKAAEKVIELEPGSELAKEAKRIIEKAAKMLADILRKEMEKIREETEEVKKEIEESKKRPQSE
SAKNLILIMQLLINQIRLLALQIRMLVLQLIL
